# Secretion encoded single-cell sequencing (SEC-seq) uncovers gene expression signatures associated with high VEGF-A secretion in mesenchymal stromal cells

**DOI:** 10.1101/2023.01.07.523110

**Authors:** Shreya Udani, Justin Langerman, Doyeon Koo, Sevana Baghdasarian, Brian Cheng, Simran Kang, Citradewi Soemardy, Joseph de Rutte, Kathrin Plath, Dino Di Carlo

## Abstract

Cells secrete numerous bioactive molecules essential for the function of healthy organisms. However, there are no scalable methods to link individual cell secretions to their transcriptional state. By developing and using secretion encoded single-cell sequencing (SEC-seq), which exploits hydrogel nanovials to capture individual cells and their secretions, we simultaneously measured the secretion of vascular endothelial growth factor A (VEGF-A) and the transcriptome for thousands of individual mesenchymal stromal cells (MSCs). We found that VEGF-A secretion is heterogeneous across the cell population and lowly correlated with the *VEGFA* transcript level. While there is a modest population-wide increase in VEGF-A secretion by hypoxic induction, highest VEGF-A secretion across normoxic and hypoxic culture conditions occurs in a subpopulation of MSCs characterized by a unique gene expression signature. Taken together, SEC-seq enables the identification of specific genes involved in the control of secretory states, which may be exploited for developing means to modulate cellular secretion for disease treatment.

## Introduction

Cell function is defined by a myriad of biomolecules that they secrete. Over 3000 proteins are predicted to be secreted from human cells^1^, and secreted proteins such as immunoglobulins, cytokines, chemokines, extracellular matrix proteins, proteases, morphogens and growth factors span a diversity of critical functions^2^. For example, mesenchymal stromal cells (MSCs) have been widely evaluated as therapeutics because they secrete bioactive factors including growth and neurotrophic factors (VEGF, HGF, GDNF), cytokines (IDO, PGE2, TGF-β, IL-10), and extracellular vesicles, which promote immunomodulation and regeneration^3,4^. However, there is a lack of methods to probe the heterogeneity in secretory functions and link these to specific gene expression networks. Methods to sort therapeutic cell populations based on functional potency and uncover the single-cell level gene expression driving this potency can be transformative for the next generation of cell therapies^2,5^.

Critical questions include which cells secrete specific proteins (heterogeneity in protein secretion), whether there is coordinated secretion among proteins, and what mechanisms control secretion. Most widely used secretion assays are bulk measurements of protein production (ELISA and cytokine arrays) which obscure phenotypic heterogeneity. ELISpot allows for single-cell secretion profiling, but it is not suited to select cells of interest for single cell RNA-sequencing (scRNA-seq), which is critical to link secretion phenotypes with underlying gene circuits. Intracellular protein/cytokine staining requires cell fixation and permeabilization and may substantially degrade mRNAs. Microfluidic droplet-based encapsulation has been used to measure secretions from cells in suspension^6–9^, but is not suited for simultaneous capture of transcription information nor for the analysis of adherent cells including MSCs. Thus, tools are still lacking to simultaneously characterize both cell secretion and mRNA at the single-cell level, for both suspension and adherent cells.

Recently, we introduced a “lab on a particle” platform to sort cells based on secretory function using standard fluorescence activated cell sorting (FACS). The approach uses microscale hydrogel particles with cavities, called nanovials, to capture single cells and their secretions^10,11^. Secretions from cells within the nanovials can be detected by staining with fluorescently labeled antibodies and sorted based on fluorescent signal. This approach has been demonstrated in previous work characterizing secreted antibodies from B cells, Chinese Hamster Ovary cells^10,12^, and of cytokines from T cells^13^.

Here, we developed a new method using the nanovial platform to profile the secretion phenotype of single cells and elucidate the associated transcriptome using scRNA-seq. Secretion encoded single-cell sequencing (SEC-seq) leverages nanovials and standard scRNA-seq platforms, such as the 10X Genomics Chromium, to retain and analyze both transcriptome and secretion information from thousands of individual cells (Fig. 1). Cells are loaded into gelatin-coated nanovials conjugated with capture antibodies for a secreted protein of interest, allowing the cells to adhere to the nanovial surface and secreted protein to bind to the antibodies. After an appropriate incubation time, nanovials are stained with oligonucleotide (oligo)-barcoded detection antibodies against the secreted product. Single-cell loaded nanovial samples, enriched by FACS, are then partitioned for downstream scRNA-seq, followed by library preparation for mRNA and oligo-barcode detection, sequencing, and data analysis employing established workflows used in other barcode-based multi-omics approaches^14,15^. The SEC-seq workflow links the secretory functions of cells to their underlying transcriptomes, for the first time at scale.

**Figure 1.**
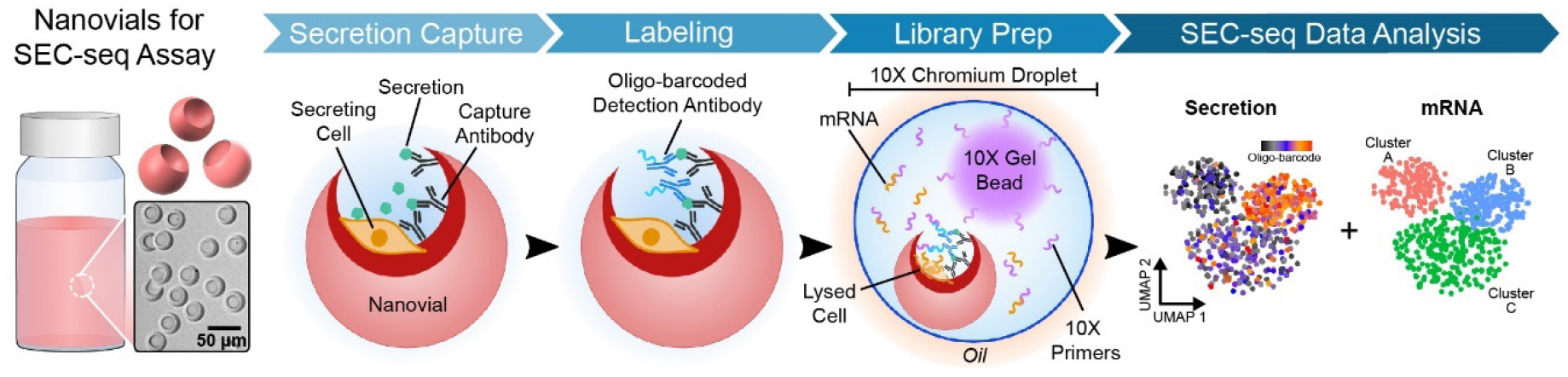
Overview of the SEC-seq workflow using nanovials. Cells are loaded, adhered, and incubated in gelatin-coated nanovials functionalized with secretion capture antibodies. Secreted and captured proteins are labeled with oligonucleotide-barcoded detection antibodies. Nanovials loaded with single cells are then introduced into the 10X Chromium workflow for library preparation. Sequencing of resulting libraries results in matched secretion and transcriptomics data for downstream analyses.

We show the utility of SEC-seq by exploring the relationship between the secretion of vascular endothelial growth factor A (VEGF-A) and the underlying transcriptome state in MSCs under normoxic and hypoxic conditions. We focused on measuring VEGF-A secretion since this growth factor is an important secreted product that promotes angiogenesis and is involved in the mechanism of action of many MSC-based therapies^16–18^. We found a relatively low correlation between VEGF-A secretion and *VEGFA* transcript levels across cells in each culture condition. Yet, a subset of cells found in both normoxic and hypoxic culture conditions secrete high levels of VEGF-A. This subpopulation is not defined by increased *VEGFA* transcript levels; instead, it is characterized by a unique, secretory gene-enriched transcription profile. We coined this gene expression signature the ‘Vascular Regenerative Signal’ (VRS) based on the presence of transcripts related to cell motility, blood vessel development, and wound response, and its link to the high VEGF-A secretory state. In addition, the induction of the hypoxic response globally elevates *VEGFA* transcript levels and VEGF-A secretion across all cells, indicating that the modulation of *VEGFA* transcript levels can tune the secretion output under certain conditions. Together, our findings suggest that multiple regulatory pathways control VEGF-A secretion, which could only be revealed by simultaneous measurement of protein secretion and the transcriptome from thousands of single cells.

Overall, our findings demonstrate the need to probe secretion and transcriptome simultaneously to better understand the regulation of protein secretion. We suggest that SEC-seq can help to develop the next generation of cell therapies by functionally defining subpopulations of cells with the most therapeutic potential and bridging function to underlying gene networks which can be engineered to improve regenerative medicine approaches.

## Results

### Establishing the SEC-seq Workflow

To develop the SEC-seq assay, we ascertained that (1) VEGF-A secretion from single MSCs can be captured on nanovials, and cells on nanovials maintain viability after sorting; (2) cell-loaded nanovials can be emulsified in a commercial microfluidic device; and (3) mRNAs can be captured and reverse transcribed from individual cells in the presence of nanovials and oligo-barcodes bound to nanovials can be quantified (Fig. 2a). We also aimed to achieve these steps using standard scRNA-seq lab equipment including a fluorescent activated cell sorter (FACS), a 10X Chromium controller for single-cell processing and library preparation, and an Illumina next-generation sequencer.

**Figure 2.**
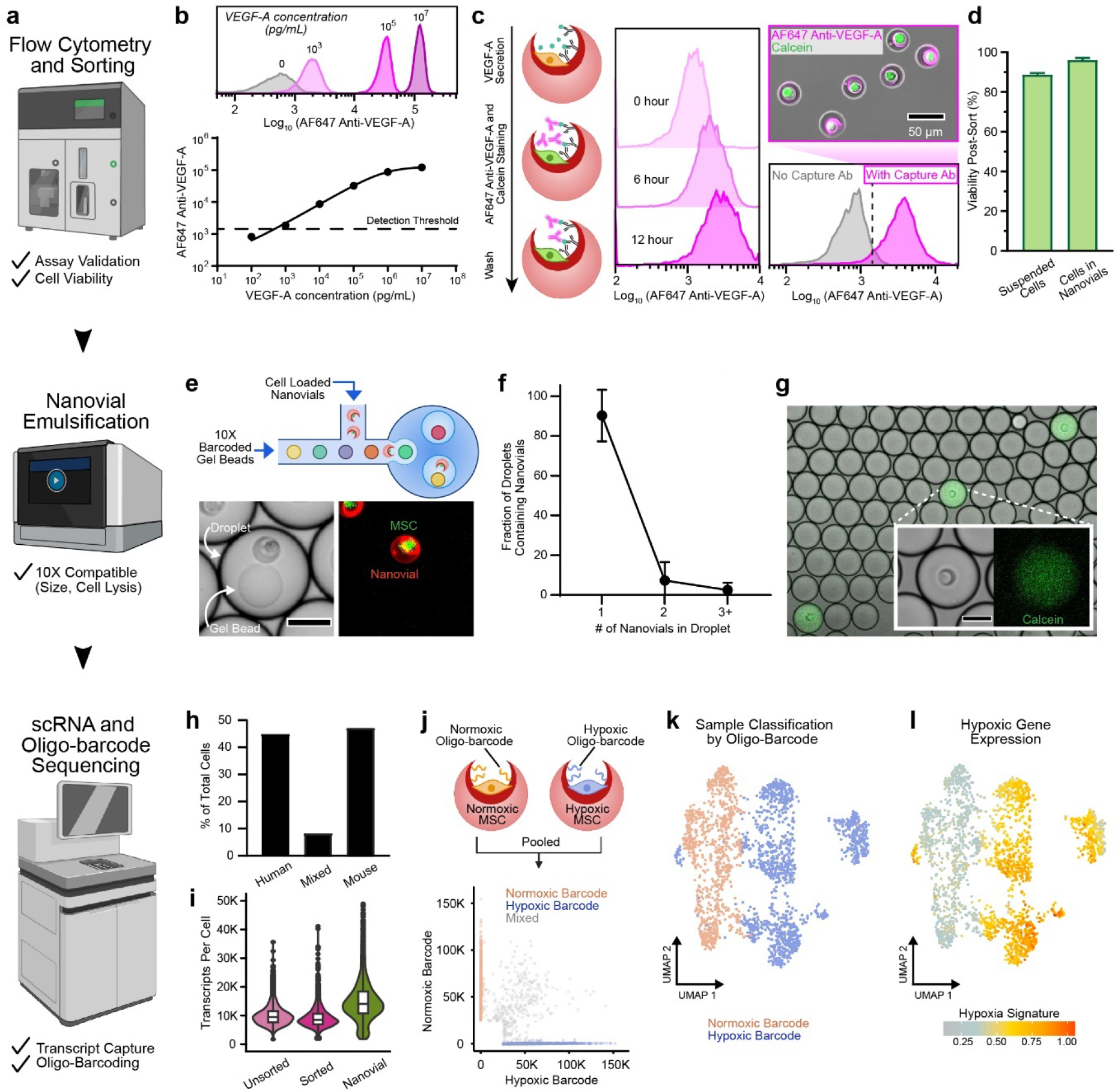
Establishing the SEC-seq workflow using nanovials. **a**, Standard scRNA-seq equipment including Sony cell sorter, 10X Chromium scRNA-seq device, and Illumina sequencers are used in the different steps of the SEC-seq workflow (top to bottom). Checkmarks indicate the steps that were validated with corresponding equipment. **b**, Standard curve of VEGF-A on nanovials using recombinant VEGF-A, immobilized via the VEGF-A capture antibody, and detected with AF647-tagged anti-VEGF-A detection antibody shows a dynamic range across 3 orders of magnitude. Horizontal line represents the detection threshold (see methods). **c**, (left) Schematic showing the steps of the VEGF-A secretion assay in single MSC-loaded nanovials. (middle) Flow cytometry histograms of VEGF-A secretion from single MSCs on nanovials after 0, 6, 12 hours of incubation. (right bottom) VEGF-A secretion assay on single MSC-loaded nanovials with and without VEGF-A capture antibody (Ab). More than 90% of cells in nanovials with capture antibody had fluorescence signal above the threshold (dotted line). (right top) The fluorescence microscopy image shows single MSCs on nanovials with secreted VEGF-A detected with a fluorescently (AF647)-tagged VEGF-A detection antibody (magenta) and cells stained with calcein AM (green). Scale bar is 50 µm. **d**, Bar plot shows cell viability measured by image analysis of live/dead stain (see methods) following flow sorting of cells in suspension or loaded in nanovials. **e**, (top) Schematic of nanovial loading into droplets with 10X training gel beads in the absence of detergent (to prevent cell lysis). (bottom) Brightfield and fluorescence images of nanovials (red) with single MSCs (green) together with a gel bead in a droplet following emulsification. Scale bar is 50 µm. **f**, Graph showing the proportion of all nanovial-containing droplets with the indicated number of nanovials. **g**, After droplet formation and in the presence of lysis buffer, lysis of calcein (green)-labeled cells on nanovials was observed by diffusion of the green fluorescent signal throughout the droplets containing single-MSC loaded nanovials. Overlaid fluorescence and brightfield images of droplets generated with a 10X Genomics NextGEM kit. Scale bar is 50 µm. **h**, Distribution of species-specific reads from a scRNA-seq experiment with nanovials containing human MSCs or mouse fibroblasts pooled in a 1:1 ratio. Species identity was called by mapping to a joined genome contig and determining the ratio of reads from each species’ genome. **i**, Comparison of transcripts per cell for either suspended (unsorted) MSCs, suspended and sorted (sorted) MSCs, or MSCs loaded on nanovials and sorted (nanovial). **j**, Scheme explaining the experiment where MSCs cultured under normoxic and hypoxic conditions, respectively, were loaded on nanovials labeled with different oligo-barcoded streptavidin molecules (‘normoxic’ and ‘hypoxic’ barcodes) and analyzed in a 1:1 ratio in a single 10X channel. The scatter plot below depicts the assignment of cells based on the normoxic or hypoxic oligo-barcode attached to nanovials via streptavidin. Mixed cells have a signal for both barcodes. **k**, UMAP plots of the scRNA-seq data derived from the experiments in (j), where each cell is labeled by their oligo-barcode assignment. Mixed cells are excluded in UMAPs. **l**, The UMAP from (k) labeled by the hypoxic gene expression signature to identify MSCs cultured in hypoxic conditions.

Specifically, we first developed a flow cytometry-compatible, single-cell VEGF-A secretion assay using nanovials. Initially, we confirmed that we can quantify VEGF-A attached to nanovials. Here, we bound increasing amounts of recombinant VEGF-A to nanovials via VEGF-A capture antibodies and measured VEGF-A retained in nanovials by flow cytometry upon incubation with fluorescently (AF647)-tagged anti-VEGF-A detection antibodies. We found that VEGF-A can be detected across a dynamic range of at least three orders of magnitude (Fig. 2b). We next asked if VEGF-A secretion from single MSCs can be detected within nanovials. MSCs were loaded into 35 µm gelatin-coated nanovials with a 20 µm cavity by simple pipet-mixing, which allows the cells to adhere to the gelatin coating via integrin binding (Fig. S1a). We found that up to 23% of nanovials contained single MSCs, which could be reliably sorted by FACS (Fig. S1b-d). To measure VEGF-A secretion from single cells, we loaded MSCs into nanovials conjugated with the VEGF-A capture antibody, and incubated them on nanovials to allow for VEGF-A secretion and binding to the capture antibody. Upon addition of the fluorescently (AF647)-tagged anti-VEGF-A detection antibody, FACS showed that nanovial fluorescence increased with the amount of time MSCs were allowed to secrete, from 0 to 12 hours (Fig. 2c, S1e). We chose a 12-hour incubation for VEGF-A secretion accumulation as it yielded a majority of the secretion signal in the observed dynamic range, with over 90% of nanovials displaying a higher signal than control nanovials with no VEGF-A capture antibody, indicating that most MSCs are secreting VEGF-A (Fig. 2c). Cell-loaded nanovials had a 2.6-fold increase in median fluorescent signal compared to empty nanovials, showing that secreted VEGF-A remained localized to nanovials containing the secreting cells with little crosstalk (Fig. S1e). Together, these experiments demonstrate the ability to load MSCs into nanovials, capture robust VEGF-A secretion from individual MSCs, and isolate MSC-loaded nanovials by FACS.

Viability is a key requirement for high quality scRNA-seq data, as it avoids cell composition bias and increases the number of cells that pass quality filtering during post-processing^19^. Since FACS is required to enrich for nanovials containing single cells, we next explored if cell viability is preserved on nanovials following FACS. We sorted suspended MSCs or nanovials containing single MSCs and found that cells in both conditions had a high viability post-sort (Fig. 2d). Additionally, nanovials protected MSCs during sorting when exposed to surfactant, suggesting that nanovials can shield cells from external stressors (Fig. S2a). Consistent with this, modeling of fluid shear stresses in the flow cytometer nozzle yielded a 400-fold higher value for fluid dynamic shear stress acting on suspended cells compared to cells loaded in a nanovial (Fig. S2b-d), suggesting that nanovials can shield cells from fluid shear stress^20^.

After establishing the nanovial-based VEGF-A secretion assay and confirming cell viability on nanovials, we explored whether cell-loaded nanovials are compatible with the emulsion generation and cell lysis required for scRNA-seq. We decided to use the 10X Chromium system as it is a commercially available scRNA-seq solution with microfluidics chip dimensions and resulting droplets that are compatible with nanovials. Accordingly, nanovials with mean diameters of 35 µm (used in the above experiments) could be successfully loaded into microfluidic droplets with barcoded primer beads (Fig. 2e). Based on image analysis of the emulsion droplets, we found that the majority of droplets with nanovials contain one nanovial, and that the inclusion of multiple nanovials is similar to the expected multiplet rate reported for cell loading by the manufacturer (9.7% vs 4%, Fig. 2f). Fluorescence microscopy of emulsions formed with nanovials containing single calcein-stained MSCs in the presence of cell lysis buffer showed localized release of the dye, pervading only nanovial-containing droplets (Fig. 2g). This result indicated successful cell lysis post droplet formation and encapsulation. Based on these findings, we concluded that cell-loaded nanovials are compatible with the 10X Chromium-based emulsification workflow.

We next explored whether scRNA-seq libraries could be successfully retrieved. Initially, we attempted a species mixing experiment, in which human MSCs and mouse fibroblasts were loaded separately into nanovials, sorted for single cells, and combined at a 1:1 ratio before loading into the 10X chip. This time, we progressed towards scRNA-seq library construction and sequencing, and retrieved transcripts from 6296 cells. The fraction of human and mouse cells reflected the initial pooling ratio with 44.8% human cells, 47% mouse cells, and 8.2% mixed cells (Fig. 2h). Importantly, these results showed that we can capture transcript data from individual cells loaded in nanovials without cell-to-cell mixing.

Next, we explored how loading MSCs in nanovials affects transcript recovery in comparison to scRNA-seq performed for free (suspended) MSCs. We performed scRNA-seq for suspended MSCs (unsorted), suspended MSCs sorted by positive calcein signal (sorted), and MSCs loaded in nanovials sorted for single cells (nanovial). Although fewer cells were detected in the nanovial sample (unsorted = 3559, sorted = 3297, nanovial = 2391), transcript detection per cell was not adversely affected and actually increased (Fig. 2i). A similar trend was observed for genes detected per cell (Fig. S3a). Both nanovial-loaded and suspended MSCs expressed standard MSC-specific surface markers (Fig. S3b); yet, cells on nanovials clustered separately from suspended cells (Fig. S3c). Gene ontology analysis showed that genes upregulated in nanovial-loaded cells are related to cell division and DNA replication (Fig. S3d,e). The higher number of unique transcripts and genes for cells in nanovials and the enhanced cell cycle signature may reflect a healthier cell state for adherent cells compared to recently dissociated MSCs. Together, these data suggest that adhered cells maintain anchorage-dependent processes like cell division^21^ and show that nanovial loading does not adversely affect transcript capture or gene expression states.

The last step in establishing the complete SEC-seq approach was to ensure accurate detection of oligo-barcodes within nanovials together with the transcriptome of single cells. To validate this step, we prepared two populations of MSCs loaded on nanovials, each population of nanovials functionalized with a specific oligo-barcoded streptavidin (Fig. 2j). The MSCs in these two nanovial samples differed as they were cultured in either standard normoxic conditions or treated with the hypoxia-mimetic agent deferoxamine (DFX), which is known to induce a hypoxic gene expression signature and to increase the secretion of angiogenic growth factors, including VEGF-A^22,23^ (Fig. S4a). Before progressing to scRNA-seq, we confirmed the increase of VEGF-A secretion in hypoxic conditions with flow cytometry for MSCs in nanovials (Fig. S4b). We expected that the hypoxia-induced gene signature would allow us to distinguish the two batches of MSCs and allow us to assess the accuracy of the oligo-barcode assignment in our pooling experiment. We found 85% of cells could successfully be assigned to one of the two oligo-barcodes and the two sets of barcoded cells separated when plotted together in a Uniform Manifold Approximation and Projection (UMAP) plot (Fig. 2j,k). The barcode-based classification matched the expected gene expression profile, as over 90% of cells with high expression of the hypoxic gene signature (which includes *VEGFA*, Supp. Table 1) were assigned to the respective (hypoxic) barcode (Fig. 2l). Using the same strategy to infer how many cells separate from barcoded nanovials between sorting and emulsion formation, we found that 93% of cells retained high levels of a single pan-sample oligo-barcode in a typical scRNA-seq experiment, indicating low cell loss from nanovials in this process (Fig. S3f). These results confirmed the ability to detect oligo-barcodes on nanovials and link barcode information to the single cell transcriptome.

### SEC-seq: Measuring VEGF-A Secretion from MSCs Reveals Uncoupled Secretion and Transcription

With all steps in the SEC-seq workflow validated (Figure 2), we next employed the SEC-seq approach for simultaneously measuring secretion of VEGF-A and the global transcriptome for thousands of individual MSCs (Fig. 3a). To detect secreted VEGF-A through a sequencing readout, we designed a custom VEGF-A detection antibody conjugated to a 10X Genomics compatible oligo-barcode (Fig. S5a). Capture primers from the gel beads add unique molecular identifiers (UMIs) and cell barcodes to this antibody-derived barcode sequence upon capture and amplification. The oligo-barcoded antibody detects VEGF-A on nanovials specifically as shown with a flow cytometry-based readout of nanovials loaded with and without recombinant VEGF-A (Fig. S5b-c).

**Figure 3.**
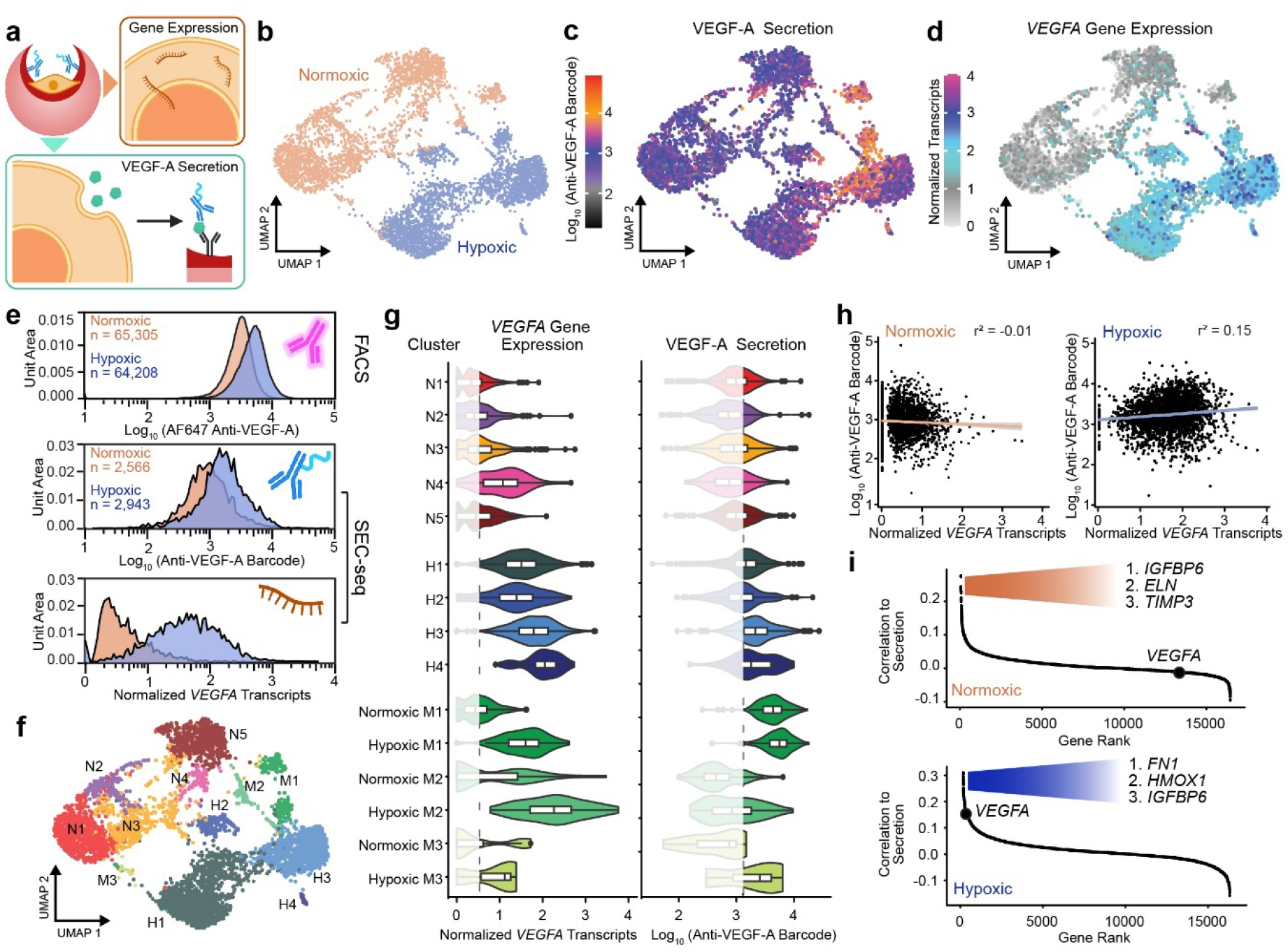
SEC-seq measuring the transcriptome and VEGF-A secretion of normoxic or hypoxic MSCs. **a**, Schematic of the detection of secreted VEGF-A protein and corresponding global gene expression for individual MSCs using the SEC-seq method. **b**, UMAP dimensionality reduction based on transcriptomes from SEC-seq experiments on normoxic and hypoxic MSCs in nanovials. Cells are labeled according to the culture condition. **c**, UMAP displaying VEGF-A secretion level, shown as log transformation of the UMI collapsed anti-VEGF-A oligo-barcode reads per cell. **d**, UMAP displaying *VEGFA* transcript levels, shown as normalized transcripts per cell. **e**, Distribution of VEGF-A secretion for cell-loaded nanovials in normoxic and hypoxic conditions, detected by FACS using a fluorescent anti-VEGF-A antibody (top) or by the SEC-seq experiment in (b) using the oligo-barcoded anti-VEGF-A antibody (middle). The last plot shows distribution of *VEGFA* transcript levels from the SEC-seq experiment cells in (b) (bottom). **f**, UMAP displaying cluster assignment. **g**, Violin plots by cluster showing *VEGFA* transcripts and VEGF-A secretion levels for all cells in the normoxic clusters (N1-N5, red shades), hypoxic clusters (H1-4, blue shades), and mixed clusters (M1-3, green shades) from (f). For mixed clusters, the levels are shown separately for normoxic and hypoxic cells. The dashed line represents the mean across all cells for each plot. Data below this threshold are lightened to highlight differences. **h**, Scatter plots showing the relationship between *VEGFA* transcript and VEGF-A secretion levels for individual normoxic (left) and hypoxic (right) cells from the experiment in (b). Best fit regression lines and Pearson correlation coefficients are shown. **i**, Plot showing the ranking of all detected genes based on the correlation of their transcript levels to the VEGF-A secretion level per cell for normoxic (top) and hypoxic (bottom) MSCs. The rank of the *VEGFA* gene is highlighted, and the top three genes per sample are also noted.

Upon validation of the oligo-barcoded VEGF-A antibody, we proceeded with the complete SEC-seq pipeline, where single MSCs were loaded in nanovials coated with VEGF-A capture antibodies and incubated for 12 hours to collect secreted VEGF-A. Nanovials were then labeled with the oligo-barcoded anti-VEGF-A detection antibody, sorted to isolate the nanovials containing single viable MSCs, and processed for scRNA-seq and oligo-barcode library preparation using the 10X workflow. This experiment was performed for MSCs in normoxic and hypoxic culture conditions and the UMI counting of oligo-barcodes per cell was used as a measure for VEGF-A secretion.

In this SEC-seq experiment, MSCs cultured in normoxic and hypoxic conditions separated in transcriptional space as depicted upon dimensionality reduction in a UMAP plot (Fig. 3b), consistent with our scRNA-seq experiment in Fig. 2j. Inspecting VEGF-A secretion and transcript levels, we made two key observations. First, we found that VEGF-A secretion is highly heterogenous across MSCs cultured in normoxic and hypoxic conditions (Fig. 3c). Although the oligo-barcoded anti-VEGF-A detection antibody was detected in nearly all normoxic and hypoxic cells, some cells in each culture conditions displayed higher VEGF-A secretion (Fig. 3c). Moreover, cells with higher VEGF-A secretion and *VEGFA* transcripts appeared to be more abundant in the hypoxic culture condition (Fig. 3c,d). On average, hypoxic cells secreted 1.72 times more VEGF-A than normoxic cells (Fig. 3e, middle). This VEGF-A secretion measurement based on SEC-seq was confirmed when the VEGF-A secretion level was determined with a fluorescently labeled anti-VEGF-A detection antibody and detection by flow cytometry (1.66 times higher in hypoxic conditions; Fig. 3e, top). Importantly, this comparison revealed that the relative magnitude and sensitivity of SEC-seq and a fluorescence-based detection assay are similar and that normoxia/hypoxia differences are within the same order of magnitude with both methods.

Second, we noted that the heterogeneity in VEGF-A secretion was not matched by similar changes in *VEGFA* transcript levels in each culture condition (compare Fig. 3c and 3d), indicating that transcript levels were less dynamic within each culture condition than secretion levels. As expected, *VEGFA* transcript levels appeared generally higher in MSCs cultured in hypoxic conditions compared to normoxic cells (Fig. 3d). However, the increase in VEGF-A secretion did not match the magnitude of change in transcript levels between the two culture conditions. Specifically, hypoxic MSCs, on average, have a 13-fold higher *VEGFA* transcript levels than normoxic cells and cover a much greater dynamic range (Fig. 3e, bottom). Together, these results suggested a surprising uncoupling between VEGF-A secretion and *VEGFA* transcript levels and an unexpected heterogeneity in VEGF-A secretion that we aimed to explore further.

To take a closer look at the relationship between VEGF-A secretion and *VEGFA* transcript levels, we divided the cells into clusters based on transcriptome (Fig. 3f). Clusters N1-N5 are predominantly populated by cells from the normoxic condition, clusters H1-H4 by cells from the hypoxic conditions, and clusters M1-M3 were formed by cells from both culture conditions (Fig S6a). The hypoxic cell clusters had higher *VEGFA* transcript and protein secretion levels than the normoxic cell clusters (Fig. 3g). However, the cluster with highest VEGF-A secretion contained cells from the normoxic and hypoxic conditions (cluster M1). Compared to all other clusters, this cluster had the highest minimum and median secretion levels. Interestingly, despite higher VEGF-A secretion in cluster M1, *VEGFA* transcript levels in this cluster were similar to other clusters (Fig. 3g). These data indicate that i) there is a subpopulation of normoxic cells (cluster M1) with increased VEGF-A secretion in the absence of higher VEGFA transcript levels; ii) the hypoxia-induced global increase in *VEGFA* transcript level is associated with a broad increase in VEGF-A secretion; and iii) cells in cluster M1 are secreting the highest level of VEGF-A compared to the clusters even in hypoxic conditions. Together, these findings confirmed the disconnect between VEGF-A secretion and transcript levels, particularly for normoxic cells, and uncovered a unique cell state associated with high VEGF-A secretion in both hypoxic and normoxic conditions.

To further examine the relationship between transcript levels and protein secretion, we calculated the correlation between *VEGFA* transcripts and VEGF-A secretion for normoxic and hypoxic cells. We found that these features were uncorrelated in normoxic cells (r^2^ = −0.01) and weakly correlated in hypoxic cells (r^2^ = 0.15; Fig. 3h). A repeat SEC-seq experiment for normoxic cells corroborated this lack of association between *VEGFA* transcripts and secretion in normoxic cells (see below, r^2^ = 0.05). When ranking all genes according to the correlation of their transcript levels to VEGF-A secretion, the *VEGFA* transcript ranked lowly in the normoxic cells (rank 13,323, Fig. 3i, Supp. Table 3). This analysis confirmed that there is no strong relationship between the basal levels of *VEGFA* transcripts and VEGF-A secretion in normoxic MSCs. In contrast, *VEGFA* was within the top 2% of correlated genes in hypoxic cells (rank 215, Fig. 3i), and when calculating transcript-to-secretion correlation across normoxic and hypoxic cells, we found a higher correlation between *VEGFA* transcripts and protein secretion (r^2^ = 0.24, Fig. S6b). Thus, the induction of the hypoxia response triggers high levels of *VEGFA* transcription which yields a modest correlation between transcription and secretion, either through a direct or secondary effect. However, overall, the *VEGFA* transcript level has surprisingly minimal input on the VEGF-A secretion amount at a single cell level.

### The MSC Subpopulation Secreting the Highest Levels of VEGF-A is Defined by a ‘Vascular Regenerative Signal’

Given that *VEGFA* transcript levels were not predictive at a single cell level of the highest secreting cells within the sample, in either normoxic or hypoxic culture conditions (Fig 3i), we were interested in defining genes that are better predictors of VEGF-A secretion levels. An examination of the top-correlating genes (Fig. 3i) demonstrated that they overlapped between normoxic and hypoxic cells (Fig. 4a). These genes included insulin growth factor binding protein 5 (*IGFBP5*) and 6 (*IGFBP6*), tissue inhibitor of metalloproteinase 3 *(TIMP3)*, heme oxygenase-1 (*HMOX1)*, fibronectin (*FN1)*, and signal peptide-CUB-EGF-like-domain-containing protein 3 (*SCUBE3*) (Fig. 4b). *IGFBP5* is a secreted signaling protein that can regulate cell growth and migration^24,25^ and *IGFBP6* was previously linked to VEGF-A secretion in a growth factor array assay for an MSC product (Stempeucel), in which VEGF-A and IGFBP6 were the top two secreted proteins in conditioned media^16^. As the transcript levels of these genes were all more highly correlated with VEGF-A secretion than *VEGFA* transcripts in both normoxic and hypoxic cells, we reasoned that they were likely to identify the most highly VEGF-A secreting cells within each cluster (Fig. 3f, g). Therefore, we inspected their expression distribution in normoxic and hypoxic cells in the SEC-seq experiment described in Figure 3 and found that cells in cluster M1 displayed the highest transcript levels for these genes (Fig. 4c, Fig. S7a).

**Figure 4.**
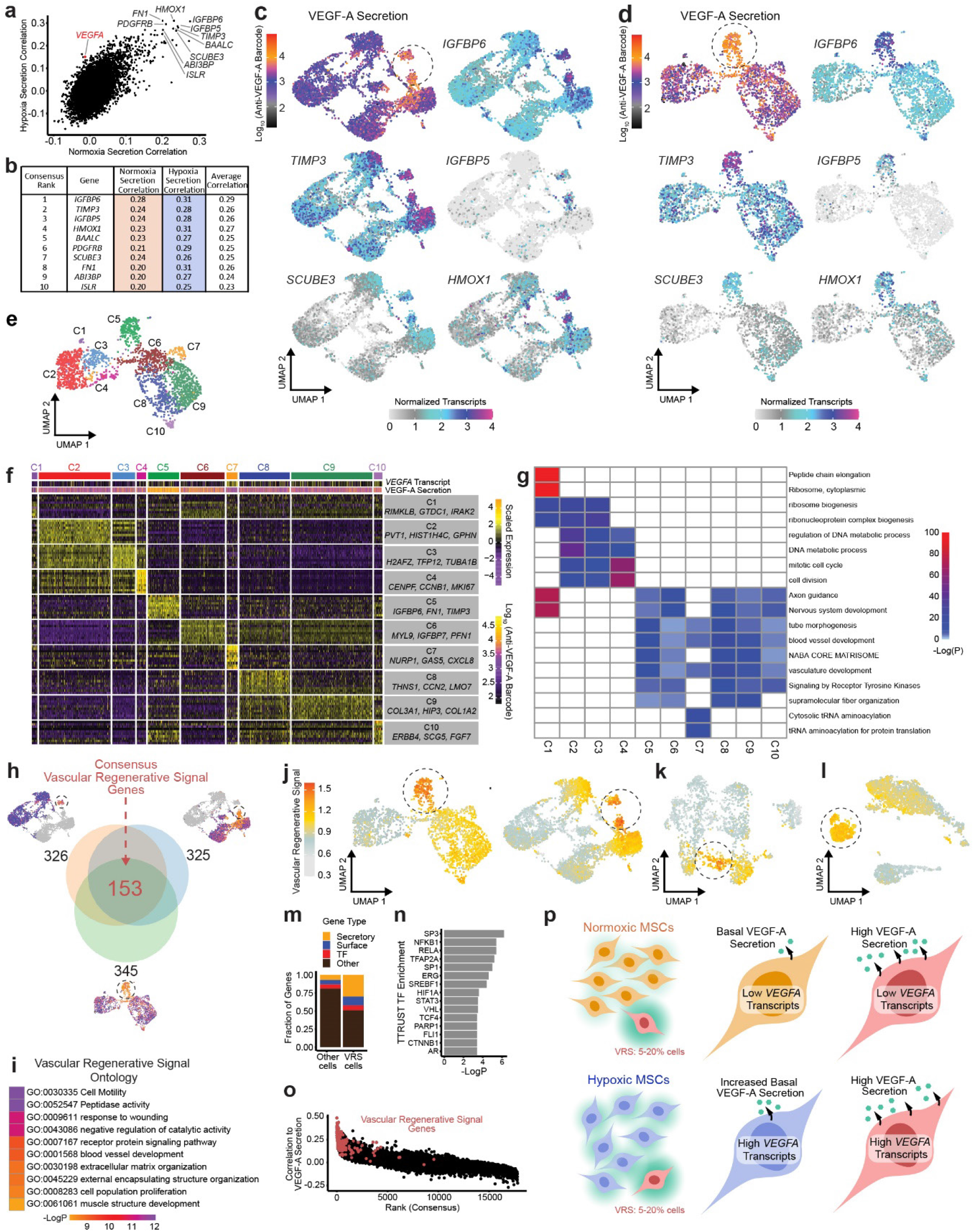
Characterization of the high-VEGF-A secreting MSC subpopulation. **a**, Scatter plot of the transcript to VEGF-A secretion correlation for all genes from SEC-seq experiments for normoxic and hypoxic MSCs from Figure 3. The 10 most highly correlating genes based on both experiments are labeled. **b**, Table giving the ranking (based on average correlation), gene name, correlation to secretion in normoxic and hypoxic cells, and the average of those two values for the ten top genes from (a). **c**, UMAPs showing VEGF-A secretion levels and expression of 5 select correlated genes from (b) in normoxic and hypoxic MSCs from Figure 3. The VEGF-A secretion UMAP is given from Fig. 3c for comparison. **d**, As in (c), for a separate SEC-seq experiment performed on MSCs in the normoxic culture condition. **e**, Cell clusters projected onto the UMAP of the replicate SEC-seq experiment **f**, Heatmap of the top 10 differentially expressed genes from each cluster (indicated on top) of the SEC-seq experiment in (c,d) (rows=genes, columns=individual cells). Displayed at the top are the log transformed VEGF-A secretion and *VEGFA* transcript levels. Right: top 3 genes differentially expressed gene for each cluster. **g**, Heatmap of the top GO terms found for all of the differentially expressed genes from the clusters in (e). The (–logP) value indicates if the term was enriched for a given cluster. **h**, Venn diagram showing the overlap of differentially expressed genes from the highly secreting cluster in 3 SEC-seq experiments (top left: normoxic MSCs from (3b), top right: hypoxic MSCs from (3b), bottom: normoxic MSCs from (4e)). Overlapping genes form the Vascular Regenerative Signal (VRS). **i**, Gene ontology analysis for VRS genes from (h). Similar terms were collapsed. **j**, Average of the normalized transcripts level of VRS genes per cell, displayed for the SEC-seq experiment in (e) and (3b). **k**, As in (j), for MSCs loaded in oligo-barcoded nanovials (see Fig. 2j-l). **l**, As in (j), for a standard scRNA-seq experiment on unsorted, suspended MSCs. **m**, Comparison of gene type classification for VRS genes and genes differentially expressed in all clusters in (e) except for those from cluster c5. **n**, Enrichment of possible TF regulators of the VRS genes based on the TRRUST database. **o**, Consensus rank of VEGF-A secretion to gene correlation based the SEC-seq experiments used in (h), with red dots displaying all VRS genes. **p**, Schematic depicting the heterogeneity of VEGF-A secretion in MSCs under normoxic and hypoxic conditions, highlighting the importance of the VRS genes for marking high VEGF-A secretion.

We validated this finding by performing and analyzing data from an additional SEC-seq experiment for normoxic MSCs (Fig. 4d). As observed previously, we found that a subpopulation of normoxic cells was marked by high VEGF-A secretion as well as high levels of *IGFBP5, IGFBP6, TIMP3*, and the rest of the top correlated genes (Fig. 4d, Fig. S7b). Clustering of the transcriptome data from this experiment revealed that the subpopulation with the highest VEGF-A secretion can be assigned to one cluster again (cluster C5; Fig. 4e, S7c). As observed before for cluster M1 (Fig. 3f,g), cluster C5 was not characterized by an increase in *VEGFA* transcript levels (Fig. S7c). Similar to the experiment described in Figure 3, *IGFBP6* was the top ranked gene when correlating transcript levels of all genes to VEGF-A secretion (r^2^=0.48) and *VEGFA* transcripts lacked correlation to secretion (r^2^=0.05; Fig. S7e-h). Furthermore, a gene expression signature formed by our previously identified 10 top-correlating genes for VEGF-A secretion (Fig. 4b) was most expressed in cluster C5, similar to the M1 cluster from Figure 3 (Fig. S7d). Thus, the steady-state transcript levels defined a set of genes reproducibly correlated with the highest VEGF-A secretion in MSCs. Taken together, these findings suggest that a subpopulation of MSCs with a defined transcriptional state, identified by a specific expression cluster in each of our SEC-seq experiments, is linked to highest VEGF-A secretion.

To further define this MSC subpopulation, we determined differentially expressed genes for each cluster for the SEC-seq experiment in Figure 4e and characterized them by gene ontology (GO) analyses (Fig. 4f,g, Supp. Table 3). Genes defining the highly VEGF-A secreting cluster C5 were associated with blood vessel morphogenesis and development, vasculature development, and extracellular matrix organization, indicative of an angiogenesis-related MSC activity and consistent with its high VEGF-A secretion state. Surprisingly, similar gene ontology terms to C5 were also found for the gene signatures defining clusters C6-C10, even though each had distinct differentially expressed genes (Fig. 4g). Due to the overlapping ontology enrichments between these clusters and lack of *VEGFA* transcript correlation, a traditional scRNA-seq analysis would have overlooked cluster C5’s identity as a super VEGF-A secretory subpopulation. The clusters with lowest VEGF-A secretion (C2, C3, and C4) were strongly enriched for mitotic signatures and likely represent actively dividing MSCs, and clusters C1 and C7 are associated with distinct but unremarkable cellular functions. Of note, the highly VEGF-A secreting cluster C5 could also be easily identified even after cell cycle regression (Fig. S8).

Since the GO analysis did not uniquely identify cluster C5 despite a unique set of highly expressed genes, we next aimed to create a gene expression signature most representative of the high VEGF-A secreting MSC subpopulation across all our SEC-seq experiments. By overlapping the marker genes expressed in the high VEGF-A secretion cluster in each of the three SEC-seq experiments (for hypoxic and normoxic MSCs in Figure 3 and normoxic MSCs in Figure 4e), we derived a consensus set of 153 genes (Fig. 4h; Supp. Table 4). GO analysis linked this gene set to extracellular matrix organization, cell motility, blood vessel development, and wound healing (Fig. 4i). On account of these GO terms and the association with highest VEGF-A secretion, we coined this shared gene set the Vascular Regenerative Signal (VRS). The VRS marked a subpopulation of cells composed of 5-20% of each sample (thresholded at 75%; see methods) in each of our MSC SEC-seq experiments (Fig. 4j), as well as in our scRNA-seq experiment testing MSCs on oligo-barcoded nanovials described in Figures 2j-l (Fig. 4k). To confirm that the VRS exists independently of nanovials and possible autocrine and/or substrate effects caused by nanovial housing of individual MSCs, we also looked for VRS-expressing cells in scRNA-seq experiments performed on free MSCs and found that 16% of cells expressed the VRS highly and that those cells were gathered within one cluster (Fig. 4l).

Interestingly, a large proportion of VRS genes classified as secretory (29%), a much larger fraction compared to the percent of secretory genes marking all other clusters (Fig. 4m), and includes secretory genes implicated in promoting angiogenesis such as tropoelastin (*ELN*), *CXCL12*, and *SCUBE3*^26–28^ (Supp. Table 4). Other functional secretory products include: HMOX1, a cryoprotective protein that maintains iron homeostasis; fibronectin 1, an extracellular matrix glycoprotein involved in tissue repair; TIMP3, an enzyme that stabilizes the extracellular matrix; and SCUBE3, a signaling protein involved in tumor angiogenesis and metastasis^29–32^. The VRS also includes some transcription factor (TF)-encoding genes known for their role in mesenchymal fate control such as *KLF6, PRRX2* and *RBPJ*^33–35^. Using a database that infers regulatory TFs for target genes based on co-mentions in publications (TRRUST^36^), we found a link between NFKB1, RELA, TFAP2A, ERG, and the hypoxia regulator HIF1A and VRS genes (Fig. 4n). As expected, VRS genes correlate highly to VEGF-A secretion, with 85% of them are within the top 500 correlating genes, as determined across all SEC-seq experiments (Fig. 4o).

Taken together, the VRS is a unique signature identifying super secretors of VEGF-A, the discovery of which was only made possible by combining the transcriptomic and secretory data of individual cells as achieved with the SEC-seq method. Our SEC-seq data demonstrate that there are multimodal transcriptional states that control VEGF-A secretion from MSCs: broadly induced cell states triggered by hypoxia as well as specialized subpopulations such as those indicated by the VRS signal (Fig. 4p).

## Discussion

Developing and using SEC-seq, we were able to link the secretion function of ∼10,000 single MSCs with their transcriptomes. Our approach greatly improves the sampling capability available compared to the previous state-of-the-art approach: splittable microwell chips in which single-cell sandwich immunoassays were performed followed by one-by-one picking of ∼20 cells for single-cell sequencing^37^. Similar to CITE-seq, which detects surface proteins together with the cell’s transcriptome using established scRNA-seq platforms such as 10x Genomics Chromium or BD Rhapsody systems^14,38^, SEC-seq uses oligo-barcoded antibodies to detect secreted proteins together with the cell’s transcriptome. Importantly, the use of nanovials enables the quantification of secreted proteins in the SEC-seq workflow. CITE-Seq has played a role in important discoveries in disease states and cell therapy development, including discovery of cell surface markers (CD80 and CD86) specific for activated regulatory T cells and a marker (CD201) for highly functional hematopoietic stem cells^39,40^. Likewise, SEC-seq should now open a new secretory dimension to multiomic studies.

SEC-seq is easily adopted by anyone who has access to current single-cell sequencing instruments, like the 10X Chromium, without the need for specialized microfluidics tools. The oligonucleotide-linked antibodies and nanovials are also accessible for others to easily repeat and build upon this work. Our approach should be extensible to probe linkages between transcriptomes and any secreted molecule for which antibodies are available and additionally enable the detection of multiple simultaneous secretions. Extending the boundaries of multiomics, SEC-seq democratizes the ability to make discoveries by linking cell secretion to gene expression (and surface markers) for thousands of cells in parallel using nanovial technology. During our development of SEC-seq, we found nanovials to provide an adhesive and shear-stress sheltering environment for cells, resulting in improved single cell analysis processes. Cell isolation schemes such as FACS sorting are often used prior to single-cell sequencing to enrich sub-populations, but many isolation or adherent cell suspension methods affect data quality. Although this work addressed secretion in adherent MSCs, suspension cells, such as B cells, are also compatible with SEC-seq through antibody capture on nanovials^41^.

MSCs have been extensively studied for cell therapy applications, leveraging an array of anti-inflammatory and pro-regenerative secreted products; yet, successful translation has been hindered by clinical outcome inconsistency attributed to functional differences that stem from the cell source and single cell level heterogeneity. Using scRNA-seq or morphological profiling, subpopulations of MSCs with various functional genotypes have been recently discovered and examined to mitigate this heterogeneity^42–44^. The primary mechanism of action for MSC therapies is thought to be through the secretion of bioactive factors that promote immunomodulation and regeneration of resident cells, such as the growth factor VEGF-A. Our data highlights the significant heterogeneity in secretion of VEGF-A in MSCs grown under both normoxic and hypoxic conditions.

Often the expression of a secretory protein is equated to secretion, however, our results suggest this assumption may not be correct across different cell types^41^. We found that *VEGFA* transcripts and VEGF-A secretion have very low correlation in normoxic and hypoxic MSCs. While surprising, the uncoupling of transcript level and protein secretion is in line with similar studies using CITE-seq where certain surface protein and transcripts feature low correlation^14,15,45^.

We uncovered the VRS as a unique transcriptomic gene signature that marks the MSC subpopulation that most highly secretes VEGF-A, which has not been previously described (Fig. 4p). Only by using SEC-seq we were able to uncover this subpopulation. The VRS contains a large fraction of secretory proteins, suggesting that SEC-seq will be valuable to further explore additional secretion products of VRS cells. Similarly, SEC-seq will be critical to functionally define the gene expression controls of the VRS and the associated VEGF-A secretion, and to understand how a particular combination of secretory genes and other regulators that, together with mesenchymal control genes, create a cell state that is permissive for high VEGF-A secretion.

The VRS could also provide a foundation for developing critical quality attributes (CQAs) for MSC therapies and the functional understanding of the link between the VRS and VEGF-A secretion could be exploited to enhance the frequency and efficacy of this potentially therapeutic population. Notably, the VRS provides genes and pathways that could be genetically targeted or modified by pre-conditioning treatments. The current standard for potency assays used as CQAs for MSC therapies relies on functional assays at the bulk level, such as production of angiogenic factors or the inhibition of T cell proliferation^46^. While this can help to exclude underperforming MSC batches, this strategy does not define a MSC subpopulation with the highest potential for therapeutic effects since it neglects the importance of cellular heterogeneity. A transcriptomic signature, such as the VRS identified in our study, that is directly tied to critical secretory functions could transform the ability to create more reproducible and reliable cell therapy products that drive clinical success for cell types like MSCs, which have high heterogeneity based on donor differences, source tissue, isolation method, and culture conditions. To this end, the VRS also contains candidate surface markers that could be more easily used for isolation if expression is correlated to protein abundance, such as *CD248, IL13RA2*, and *CDH11*.

Our finding that *VEGFA* transcription and VEGF-A protein secretion are not tightly linked in MSCs opens new questions and has important implications for engineering cell products. These questions include (1) how is VEGF-A secretion controlled and how dependent is it on cell state; (2) how are cells facilitating the build-up and release of different amounts of VEGF-A secreted protein outside of gene regulation and (3) why would the uncoupling of VEGF-A secretion and transcript levels be beneficial for ultimate biological function compared to direct control of secretion via the regulation of *VEGFA* steady state transcript levels. From a biotechnology perspective, our results suggest that approaches to drive secretion to modulate engineered cell therapy function^47^ should not rely alone on introducing copies to increase the transcript level of the target gene, but rather could benefit from a holistic engineering of the pathways involved in driving secretion, e.g. modulating the VRS in MSCs.

## Author Contributions

S.U., J.L., J.D., K.P., and D.D. conceived the idea and contributed to the design of experiments. S.U., J.L., D.K., S.B., B.C., S.K., and C.S. performed the experiments/simulations and contributed to data analysis. S.U., J.L., K.P., and D.D. drafted the manuscript, and all the authors provided feedback. S.U. and J.L. contributed equally and have the right to list their name first in their CVs.

## Methods

### List of reagents and resources

| **REAGENT OR RESOURCE** | **SOURCE** | **IDENTIFIER** |
| --- | --- | --- |
| **Antibodies** |  |  |
| Biotin anti-human VEGF Antibody (clone: Poly5225) | Biolegend | 522503 |
| Anti-VEGF Antibody, Alexa Fluor® 647 Conjugate | Sigma-Aldrich | ABS82-AF647 |
| TotalSeq-B 5136 anti-human VEGF Antibody (clone: A15136H) | Biolegend | n/a; custom antibody conjugation |
| Alexa Fluor 647 AffiniPure F(ab')<sub>2</sub> Fragment Goat Anti-Mouse IgG, F(ab')<sub>2</sub> fragment specific | Jackson ImmunoResearch | 115-606-072 |
| **Chemicals, Peptides and Recombinant Proteins** |  |  |
| *Nanovial fabrication and modification* |  |  |
| 4-arm Poly(ethylene glycol) acrylate, MW: 5kDa | Advanced BioChemicals | 4AP0901-1g |
| Gelatin from cold water fish skin | Sigma-Aldrich | G7041-100G |
| Lithium phenyl-2,4,6-trimethylbenzoylphosphinate | Sigma-Aldrich | 900889-1G |
| Sulfo-NHS-Biotin | ApexBio | A8001 |
| Hexane, mixture of isomers | Sigma-Aldrich | 178918-2.5L |
| 1H,1H,2H,2H-Perfluoro-1-octanol | Sigma-Aldrich | 370533-5G |
| Pico-Surf(5%(w/w)) in Novec™ 7500 | Sphere Fluidics | CO24 |
| 3M Novec™ 7500 | Gallade Chemical | N17501 |
| *Cell Culture and Hypoxia Inducers* |  |  |
| Mesenchymal Stem Cell Basal Medium for Adipose, Umbilical and Bone Marrow-derived MSCs | ATCC | PCS-500-030 |
| Mesenchymal Stem Cell Growth Kit for Adipose and Umbilical-derived MSCs - Low Serum | ATCC | PCS-500-040 |
| Fetal Bovine Serum US Certified | Invitrogen | 16000044 |
| Antibiotic-Antimycotic | Thermo Fisher | 15240096 |
| DMEM, high glucose, pyruvate | Invitrogen | 11995065 |
| TrypLE | Thermo Fisher | 12-604-013 |
| Deferoxamine mesylate salt | Sigma-Aldrich | D9533-1G |
| Cobalt(II) chloride hexahydrate | Sigma-Aldrich | C8661-25G |
| *Nanovial Assays* |  |  |
| Streptavidin | Thermo Fisher | 434302 |
| TotalSeq-B0952 PE Streptavidin | Biolegend | 405287 |
| TotalSeq-B0953 PE Streptavidin | Biolegend | 405289 |
| Recombinant Human VEGF 165 Protein | R&D Systems | 293-VE |
| Calcein AM | Thermo Fisher | C3099 |
| Propidium Iodide | Thermo Fisher | P3566 |
| CallTracker Orange CMTMR | Thermo Fisher | C2927 |
| DPBS (Ca<sup>+</sup>,Mg<sup>+</sup>) | Invitrogen | 14040133 |
| Bovine Serum Albumin | Sigma-Aldrich | A7906-100G |
| Pluronic F-127 | Sigma | P2443-250G |
| CellTrics Filters, 20 µm | Thermo Fisher | NC9699018 |
| Reversible Strainers (37 µm) | STEMCELL Technologies | 27215 |
| Commercial Assays |  |  |
| ELISA MAX Deluxe Set Human VEGF | Biolegend | 446504 |
| Experimental Models: Cell Lines |  |  |
| Immortalized adipose-derived MSCs | ATCC | SCRC-4000 |
| scRNA-seq and SEC-seq library preparation and sequencing |  |  |
| Chromium Next GEM Single Cell 3' Kit v3.1 | 10x Genomics | 1000268 |
| Chromium Next GEM Chip G Single Cell Kit | 10x Genomics | 1000120 |
| Dual Index Kit NN Set A | 10x Genomics | 1000243 |
| Dual Index Kit TT Set A | 10x Genomics | 1000215 |
| 3' Feature Barcode Kit | 10x Genomics | 1000262 |
| D5000 screentape and reagents | Agilent | 5067-5588, 5067-5589 |
| D1000 screentape and reagents | Agilent | 5067-5582, 5067-5583 |
| NovaSeq 6000 S4 Reagent Kit v1.5 (200 cycles) | Illumina | 20028313 |
| Software and Algorithms |  |  |
| ImageJ | NIH | https://imagej.nih.gov/ij |
| Matlab | Mathworks | Version 2021b |
| FlowJo | FlowJo, LLC | Version 10.8 |
| COMSOL Multiphysics | COMSOL | Version 6.0 |
| R | The R Foundation | Version 4.04 |
| Seurat | Satija Lab | Version 4.10 |

#### MSC Cell Culture

Immortalized human adipose-derived MSCs (ATCC SCRC-4000) were cultured in MSC Basal Medium (ATCC PCS-500-030) supplemented with Low Serum MSC Growth Kit for adipose MSCs (ATCC PCS-500-040) and Antibiotic-Antimycotic (A-A; Invitrogen) resulting in final complete MSC media concentrations of 2% Fetal Bovine Serum (FBS), 5 ng/mL FGF-1, 5 ng/mL FGF-2, 5 ng/mL EGF, 2.4 mM L-Alanyl-L-Glutamine and 1% A-A. MSCs were cultured in incubators at 37°C and 5% CO_2_ and passaged once 70-80% confluent, with MSCs up to passage 25 used in experiments.

#### MEF Cell Culture (for species mixing experiment)

Primary mouse embryonic fibroblasts (MEFs) were cultured in Dulbecco’s Modified Eagle Medium (Gibco) supplemented with 10% FBS and 1% A-A. MEFs were cultured in incubators at 37°C and 5% CO_2_ and passaged once 70-80% confluent, with MEFs up to passage 7 used in experiments.

#### VEGF ELISA on MSC Conditioned Media

To measure VEGF-A secretion from MSCs grown in tissue culture plates, we seeded 100,000 MSCs in 1.33 mL culture media in each well of 6-well plates. After 3 days, media was exchanged for normoxic or hypoxic media. Specifically, we used cell culture media for the normoxic condition and media supplemented with deferoxamine (500 µM and 1000 µM) or cobalt chloride (50 µM and 100 µM) for hypoxic conditions. After 24 hours, the conditioned media was collected and filtered, and cell number at the time of media collection was counted. The VEGF-A secretion amount was measured using a VEGF-A Human ELISA kit (Biolegend) by plate reader. The experiment was performed in triplicates for each tested condition. Secretion amount was calculated by first normalizing each sample by cell number, and then averaging the triplicate experiments.

#### Nanovial fabrication and modifications

35 µm nanovials were fabricated using a three-inlet flow-focusing microfluidic device formed from polydimethylsiloxane (PDMS). PEG pre-polymer, gelatin and oil phases were infused at flow rates of 1.5 μl/min, 1.5 μl/min and 15 μl/min, respectively. The PEG pre-polymer phase comprised 27.5% w/v 5 kDa 4-arm PEG acrylate (Advanced BioChemicals) with 4% w/v lithium phenyl-2,4,6-trimethylbenzoylphosphinate (LAP, Sigma) in phosphate buffered saline (PBS, pH 7.2). The gelatin phase comprised 20% w/v cold water fish gelatin (Sigma) in deionized water. The oil phase comprised Novec 7500 (3M) with 0.5% v/v Pico-surf (Sphere Fluidics). Oil partitioned the aqueous phases into monodisperse water-in-oil droplets and the PEG and gelatin polymers phase separated into a PEG-rich and gelatin-rich phase. Phase-separated droplets were crosslinked with focused UV light through a DAPI filter set and 10X microscope objective (Nikon, Eclipse Ti-S) near the outlet region of the microfluidic device. Polymerized nanovials were collected and any unreacted phases including oil were removed through a series of washing steps as previously described^10,12^. Biotinylation of the gelatin-layer formed in the nanovial cavity was conducted by incubating nanovials with 10 mM Sulfo-NHS-Biotin (APExBIO) overnight at room temperature while mixing. Nanovials were then washed in pluronic buffer consisting of 0.05% Pluronic F-127 (Sigma), 1% antibiotic-antimycotic (Thermo Fisher) in PBS and sterilized in 70% ethanol overnight. Sterile nanovials were stored 5x diluted (i.e. 100 μL of concentrated nanovial volume was resuspended in 400 μL pluronic buffer resulting in 6.5 million nanovials/mL) in this pluronic buffer at 4°C.

#### Conjugating VEGF-A capture antibodies to nanovials

5x diluted nanovial suspension was incubated with an equal volume of 260 μg/mL of streptavidin for 30 minutes at room temperature on a tube rotator. Excess streptavidin was washed out by pelleting nanovials at 200g for 3-5 minutes, removing supernatant and adding 1 mL of wash buffer (pluronic buffer + 0.5% bovine serum albumin [BSA]) three times. All subsequent wash steps in assay workflows were performed similarly unless otherwise noted. The 5x diluted streptavidin-coated nanovial suspension was then incubated with an equal volume of 71.5 μg/mL of biotinylated anti-human VEGF antibody (Biolegend) for 30 minutes at room temperature on a tube rotator and washed. Nanovials were resuspended at a 5x dilution in wash buffer or cell culture medium prior to the next workflow step.

#### Loading cells in nanovials

MSCs were loaded into nanovials by pipet-mixing nanovials and the cell solution in 5 ml round-bottom polypropylene tubes (Corning Falcon), which provide a cell-adhesion resistant material and vent cap for gas exchange during incubation. First, the 5x diluted nanovial suspension was reconstituted in complete MSC media. MSCs were detached from tissue culture flasks using TrypLE (Gibco) and resuspended in media at concentrations of 0.9, 1.5 and 2.2 million per ml. Cells and nanovials are then pipette-mixed 10 times in a 3:1 volume ratio for final cell to nanovial ratios of 0.4:1, 0.7:1, and 1:1, respectively. A total volume of less than 0.5 ml per tube was maintained throughout to reduce cell clumping during incubation. The nanovial suspensions were incubated at 37°C and 5% CO_2_ for 2 hours to allow MSCs to bind to the gelatin coating in nanovial cavities. Then, nanovial suspensions were first strained through a 20 µm strainer (CellTrics) to remove unbound cells. Subsequently, recovered nanovials were strained through a 37 µm strainer (STEMCELL) to remove any large cell/nanovial aggregates. The 1:1 cell to nanovial ratio was used for remaining experiments.

#### Single-cell VEGF-A secretion on nanovials

VEGF-A capture antibody-conjugated nanovials were loaded with MSCs as described above. After straining, nanovials were incubated in 6-well plates for 12 hours to allow for secretion, with up to 170,000 nanovials per well in 2 mL media (for time sweep secretion experiment, 0, 6, and 12 hours incubations were performed). Plates were shaken in horizontal cross-movements before placing in the incubator to space out nanovials. After incubation, nanovials were collected in wash buffer, centrifuged, and resuspended as 5x diluted nanovial suspension. This suspension was incubated with an equal volume of 71.5 μg/mL AF647 anti-human VEGF antibody (Sigma) for 30 minutes at 37°C with gentle vortexing every 10 minutes before washing excess antibody with one high-dilution wash (>100x nanovial volume). Samples were then analyzed by imaging using fluorescence microscopy and analyzed/sorted by FACS as described below. For the hypoxia inducer concentration sweep, MSCs in nanovials were incubated in cell culture media for normoxic condition and media was supplemented with deferoxamine (100, 250, 500, 1000 µM) or cobalt chloride (50 µM and 100 µM) for hypoxic conditions. At least 10,000 single cell-loaded nanovials were analyzed per condition.

#### Flow cytometry and sorting of nanovials using SONY SH800S

Nanovial samples were diluted 25x in washing buffer before sorting on the SONY SH800S cell sorter using a 130 µm sorting chip. The sorter featured violet (405 nm), blue (488 nm), yellow (561 nm) and red (640 nm) lasers and 450/50 nm (FL1), 525/50 nm (FL2), 600/60 nm (FL3) and 665/30 nm (FL4) filters were used. Typical sensor gain settings used for nanovial samples are given in Table 1, along with stains or antibodies measured using the filters. Compensation was performed for samples with spectral overlap (e.g. calcein spillover into FL4 for samples labeled with calcein and AF647 anti-VEGF) using an unstained control (plain nanovials) and single-stained controls (calcein-stained cells on nanovials; AF647 anti-VEGF-A-labeled recombinant VEGF in empty nanovials). The typical gating strategy for nanovials is shown in Fig. S1e. Flow cytometry data was analyzed using FlowJo software version 10.8 (BD).

**Table 1.** Typical sensor gain settings used with SONY SH800S

| Sensor | FSC | BSC | FL2 | FL3 | FL4 |
| --- | --- | --- | --- | --- | --- |
| Gain | 2 | 25% | 20% | 25% | 32% |
| Stain/Antibody |  |  | Calcein AM | CellTracker Orange | AF647 Anti-VEGF,<br>AF647 Anti-Mouse IgG |

#### Viability measurements for suspended cells and cells on nanovials after sorting

MSCs stained with CellTracker Orange CMTMR were either prepared in suspension or loaded in 35 µm nanovials. This stain was chosen to avoid spectral overlap with later live/dead Incucyte imaging. For MSCs in suspension, FACS buffer (PBS without calcium and magnesium + 1% A-A + 0.5% BSA) or FACS buffer with 0.05% Pluronic was used. For MSCs in nanovials, wash buffer (PBS with calcium and magnesium + 1% A-A + 0.5% BSA + 0.05% Pluronic) or wash buffer without 0.05% Pluronic was used. Immediately after straining the sample as described previously, samples were sorted. The 405 nm, 488 nm, and 561 nm lasers were turned on for this experiment. Samples were sorted per well in a 96 black well plate in triplicate, and a similar amount of unsorted sample was prepared. Then calcein AM and propidium iodide were added to the existing media plus sorted sample resulting in a final 1:1000 dilution of each stain, and incubated for 30 minutes before imaging on the IncuCyte Live-Cell Analysis System using phase, green and red channels. Fluorescent images were thresholded manually, and the number of live/dead cells were quantified using the IncuCyte S3 software.

#### Finite element analysis for shear stress calculations on cells in nanovials versus suspended cells

Shear stress experienced by cells in nanovials and suspended cells were modeled using a 2D axisymmetric geometry with COMSOL Multiphysics Single-Phase Laminar Flow physics. The full Navier-Stokes equations are solved by treating either the cell in nanovial or suspended cell as stationary in the center of the channel. This models the fluid flow at the sheath flow junction where the slower sample stream meets the much more rapidly flowing sheath fluid. The channel is set to a diameter of 780 µm, as measured on the junction of the SONY SH800 130 µm chip. The sample inlet boundary condition is set as fully-developed laminar flow with flow rate of 10 µL/min (sample pressure level 4), and the sheath inlet boundary condition has a flow rate of 5.5 mL/min (as measured experimentally). The outlet boundary condition is set to 0 atm pressure. The shear stress is then calculated along the surface of the suspended cell or cell within a nanovial.

#### Dynamic range of VEGF-A immunoassay on nanovials using flow cytometry

Nanovials were conjugated with biotinylated anti-VEGF-A capture antibody as mentioned earlier. 5x diluted nanovials were incubated with equal volumes of 0, 0.1, 1, 10, 100, 1000, 10000 ng/mL of recombinant human VEGF-A (R&D Systems) for 12 hours at 4°C on a tube rotator. Excess recombinant VEGF-A was washed before reconstituting as 5x dilution of nanovials and incubating with equal volume of 71.5 μg/mL AF647 anti-VEGF-A (Biolegend) at room temperature for 30 minutes in the dark on a tube rotator. After washing, nanovials were reconstituted as 25x diluted suspension in wash buffer and transferred to a flow tube. Additionally, a small fraction of the sample was transferred to a 96-well plate to be imaged on a fluorescence microscope. Fluorescent signal on nanovials was analyzed using the Sony sorter by gating for single particles. To generate the standard curve, the median signal from each concentration of VEGF-A was calculated on FlowJo and plotted against concentration. The threshold was calculated as median (0 ng/mL sample) + 3 * standard deviation (0 ng/ml sample).

#### Oligo-barcoded anti-VEGF-A immunoassay validation

To validate the oligo-barcoded anti-VEGF-A specificity on nanovials, we prepared nanovials either with or without recombinant VEGF-A bound (as discussed above, using 1000 ng/ml VEGF-A concentration). Then we incubated 5x diluted nanovials with an equal volume of either 0 or 71.5 ng/ml oligo-barcoded anti-VEGF-A (Biolegend) for 30 minutes at room temperature on a tube rotator. The four resulting samples were as follows: 1) VEGF-A^+^, anti-VEGF-A^+^, 2) VEGF-A^-^, anti-VEGF-A^+^, 3) VEGF-A^+^, anti-VEGF-A^-^, 4) VEGF-A^-^, anti-VEGF-A^-^. To measure the binding of the oligo-barcoded anti-VEGF-A detection antibodies, we then incubated the four samples with an equal volume of 71.5 ng/ml of AF647 goat anti-mouse IgG (Jackson ImmunoResearch), which binds to the oligo-barcoded Anti-VEGF-A antibody which is mouse species, for 30 minutes at room temperature on a tube rotator. Samples were then analyzed on the Sony sorter by gating for single particles.

#### Single-cell gene expression and feature barcode detection library generation for cell-loaded nanovials

We followed the standard protocol for the Chromium Next GEM Single Cell 3’ Kit v3.1 unless otherwise noted in the methods (10X Genomics, https://www.10xgenomics.com/products/single-cell/). Single cell containing nanovials were isolated using FACS, as noted above. Nanovials were prepared in PBS + 0.04% BSA at a typical concentration of 500 nanovials per µL (experiments in Fig. 2h,j were done at higher concentrations of 2k per µL). We loaded 16.5 µL of this solution containing approximately 8,250 nanovials (target cell recovery ∼ 5,000 cells) into Chip G microfluidic emulsion devices. As nanovials settle more quickly than cells, we adjusted the loading procedure to reduce the time in between sample loading and emulsion generation. During the sample loading step, the nanovial + master mix suspension was first allowed to settle for 2 minutes after mixing, then the first half (35 µl) was taken from the supernatant and loaded in the 10x chip sample well (Row 1). The gel bead well and partitioning oil wells were then loaded. The second half of the nanovial sample (35 µl) was pipet-mixed and added to row 1 immediately before loading into a Chromium X controller to generate emulsions.

Libraries were assembled from emulsions according to the protocol (Chromium Single Cell 3’ Reagent Kits User Guide (v3.1 Chemistry Dual Index) with Feature Barcoding technology for Cell Surface Protein and Cell Multiplexing), using the Chromium Single-Cell 3’ Library Kit (10X Genomics) for purification, amplification, fragmentation, end repair, A-tailing, adapter ligation, and final library indexing and amplification. Library cleanup was done with SPRI select reagent beads (Beckman Coulter, B23317). Separate libraries were made to detect the nanovial-associated streptavidin oligonucleotide barcode DNA and SEC-seq antibody-conjugated oligonucleotide DNA per cell using the protocol for Cell Surface Protein libraries in the above user guide. All libraries were quantified on a Tapestation 4200 (Agilent, G2991BA) according to the manufacturer’s protocols using the D5000 screentape and reagents for cDNA QC (Agilent, 5067-5588, 5067-5589) and D1000 screentape and reagents for final library QC (Agilent, 5067-5582, 5067-5583). Libraries were pooled and paired-end sequenced at 100bp per end with an additional 10bp of index reads on a NovaSeq S4 flow cell (Illumina, Novaseq 6000).

#### Emulsion imaging

Reservoir devices with a height of 130 µm made using standard soft lithography techniques (PDMS devices plasma bonded to glass slides) were used to image nanovial-containing emulsions produced from the 10X Chromium system in a single layer. 5 µl of 10X partitioning oil was added first to displace air, followed by 3-5 µl of 10X emulsion added gently without disrupting the droplets. To quantify nanovial loading in droplets, the emulsion was imaged using a fluorescence microscope. Then a custom MATLAB script was used to first detect all droplets, and then determine how many nanovials were in each detected droplet.

#### Species mixing experiment

Nanovials were separately loaded with either human MSCs (1 cell : 1 nanovial) or MEFs (0.4 cell : 1 nanovial). Single-cell loaded nanovials for each species were separately sorted and then combined in a 1:1 ratio. The sample was reconstituted as 2000 nanovials/µl before loading into a 10X chip for single-cell sequencing library preparation.

#### Comparing transcriptomes of suspended cells and cells on nanovials

MSCs were prepared either suspended or loaded in nanovials. The nanovial sample was incubated for 12 hours, similar to secretion assay experiments, and labeled with oligo-barcoded streptavidin (Biolegend) after loading. Suspended cells and nanovial samples were sorted for single cells or single cells on nanovials, respectively. An additional sample of suspended cells was left unsorted. Each of the three samples (suspended and sorted, suspended and unsorted, and cells on nanovials) was reconstituted as 500 cells/µl or nanovials/µl and loaded in separate 10X chip channels for single-cell sequencing library preparation.

#### Barcoded normoxic and hypoxic conditioned MSCs on nanovials pooled for scRNA-seq

MSCs were loaded into nanovials and were incubated for 12 hours in either normoxic or hypoxic (500 µM deferoxamine) media. After incubation, each nanovial sample was conjugated with a different oligo-barcoded streptavidin (Biolegend). Nanovial samples were then sorted using the sorting gate for single cells on nanovials, and then normoxic and hypoxic cell-loaded nanovials were combined in a 1:1 ratio. The pooled sample was reconstituted as 1000 nanovials/µl before loading into a 10X chip for single-cell sequencing library preparation.

#### SEC-seq for MSC transcriptome and VEGF-A secretion

VEGF-A capture antibody-conjugated nanovials were loaded with MSCs as described above. After straining, nanovials were incubated in 6-well plates for 12 hours to accumulate VEGF-A secretions, with up to 170,000 nanovials per well in 2 mL media (cell culture media for the normoxic condition and media supplemented with 500 µM deferoxamine for the hypoxia-inducing condition). Plates were shaken in horizontal cross-movements before placing in the incubator to space out nanovials. After incubation, nanovials were collected in wash buffer, centrifuged, and resuspended as 5x diluted nanovial suspension. This suspension was incubated with an equal volume of 71.5 μg/mL of oligo-barcoded anti-VEGF-A detection antibody (Biolegend) for 30 minutes at 37°C with gentle vortexing every 10 minutes before washing excess antibody with one high-dilution wash (>100x nanovial volume). Samples were then sorted using the SONY SH800S using the single cell on nanovial gate. The normoxic and hypoxic MSC on nanovial samples were then reconstituted separately as 500 nanovials/µl before loading into a 10X chip for single-cell sequencing library preparation.

### Data analysis

Sequencing data were demultiplexed in Basespace and mapped, barcode collapsed, and filtered in the Cell Ranger software (10x Genomics). Reads were mapped to the hg38 Refseq human reference transcriptome, or for the mixed species experiment, a fusion of the hg38 genome and the mouse mm38 genome build using the Cell Ranger mkref function. The output from Cell Ranger, a raw sparse matrix with digital expression of cell barcodes by genes, was used for downstream analysis.

Using Seurat 4.1 in R (https://github.com/satijalab/seurat), we performed normalization of transcripts and clustering of cells, and obtained reduced dimensionality PCA and UMAP coordinates for each cell. Cells were regressed using depth as the variable. To identify and remove doublets, the cell matrix was processed by DoubletFinder (https://github.com/chris-mcginnis-ucsf/DoubletFinder).

We used streptavidin barcodes linked to nanovials to separate mixed hypoxic and normoxic MSCs in Fig. 2. Barcode reads for this feature were matched to each cell using Cell Ranger’s multi-config workflow. We added a pseudocount to each barcode read and, for each cell, calculated the ratio of barcodes. Cells with a ratio favoring one barcode at least 2.5 fold were called for that tagged sample (“Normoxic” or Hypoxic”), while cells with ratios between 2.5 and −2.5 fold were considered “Mixed” and removed from the analysis. Cells with less than 25k streptavidin reads for either sample (∼15% of max) were considered unannotated and also discarded. In Fig. S3, we identified cell separation from nanovials during emulsion formation using detection of a single barcode; here escaped cells were called which had fewer than 800 barcode reads (25% of average read number).

Analysis and plots were mostly created using R and the libraries ggplot2 and pheatmap. The hypoxic gene signature shown in Fig. 2l was derived de novo using the gene-gene correlation method GEND, as previously described^48^. Cell clusters were derived using Seurat’s FindNeighbors and FindClusters function, and differential expression among clusters was determined by using the FindMarkers function. Differential expression between samples was determined by testing each gene for a distribution p-value of less than 0.05 and a fold change greater than 2 between the average gene value in the cells for each sample. Histograms for sequencing data was generated by grouping normalized reads into 100 equally sized bins. Gene ontology for various gene lists was determined using Metascape.

Cell cycle regression was performed using Seurat. Briefly, cell cycle genes marking S phase and G2/M phases were used to create cell cycle scores for each cell using the CellCycleScores function. These scores were used to adjust the gene by cell matrix using the ScaleData function to regress out the effect of cell cycle, and then the output was processed and analyzed using normal parameters.

SEC-seq reads were recovered from the Cell Surface Protein library workflow and matched to the transcriptome cell barcodes using Cell Ranger’s multi-config tag workflow. Secretion reads were log-transformed to more closely match the dynamic range of the normalized gene transcripts. Gene correlates were determined using Pearson’s correlation of all gene transcripts against the log of secretion reads. Secretion correlate genes were ranked by order of correlation. For generation of a consensus ordering between multiple SEC-seq experiments, the Pearson’s correlation values were averaged for each gene and a new ordering rank was determined from that average.

The Vascular Regenerative Signal (VRS) was determined by identifying the cluster with high *IGFBP6* expression in three MSC scRNA-seq experiments (two Normoxic replicates and a Hypoxic MSC run), running differential gene expression analysis for the respective cluster against all other cells in the given experiment, and then taking the overlap of genes between the three samples. The percent of cells in each experiment called as “VRS expressing” was calculated by averaging the expression of all genes in the VRS per cell and using a threshold of 75% of the max averaged value. Potential regulators of the VRS genes were determined by testing the VRS genes for enrichments in the TRRUST database^36^. VRS gene types were annotated using separate databases for the secretome(SPRomeDB)^49^, transcription factors (ATFDB)^50^, and surfactome (SURFY)^51^, after overlaps between the secretome and the other two databases were pruned.

