## Supplementary Information for "Secretion encoded single-cell sequencing (SEC-seq) uncovers gene expression signatures associated with high VEGF-A secretion in mesenchymal stromal cells"

**Table of Contents**

Figure S1. Cell loading into nanovials, enrichment of single cell-loaded nanovials by FACS, and VEGF-A nanovial secretion assay validation

Figure S2. Nanovials protect viability of MSCs during flow sorting

Figure S3. Transcript changes related to cell loading and adhesion in nanovials

Figure S4. Effect of hypoxia inducers on VEGF-A secretion by MSCs

Figure S5: Oligo-barcoded Anti-VEGF-A binding specificity on nanovials

Figure S6. Analysis of the SEC-seq experiments for normoxic and hypoxic MSCs

Figure S7. Identification of a high-VEGF-A secreting MSC subpopulation in a replicate experiment.

Figure S8. The high VEGF-A secretion cluster is not affected by cell cycle regression

Table 1: List of Hypoxic Signature Genes dervied from Hypoxic MSCs

Table 2: Correlation value and rank of the transcript levels for all detected genes relative to VEGF-A secretion

Table 3: List of genes marking the clusters in Figure 4f

Table 4: List of VRS genes and their classification as secretory protein, surface protein, or transcription factor

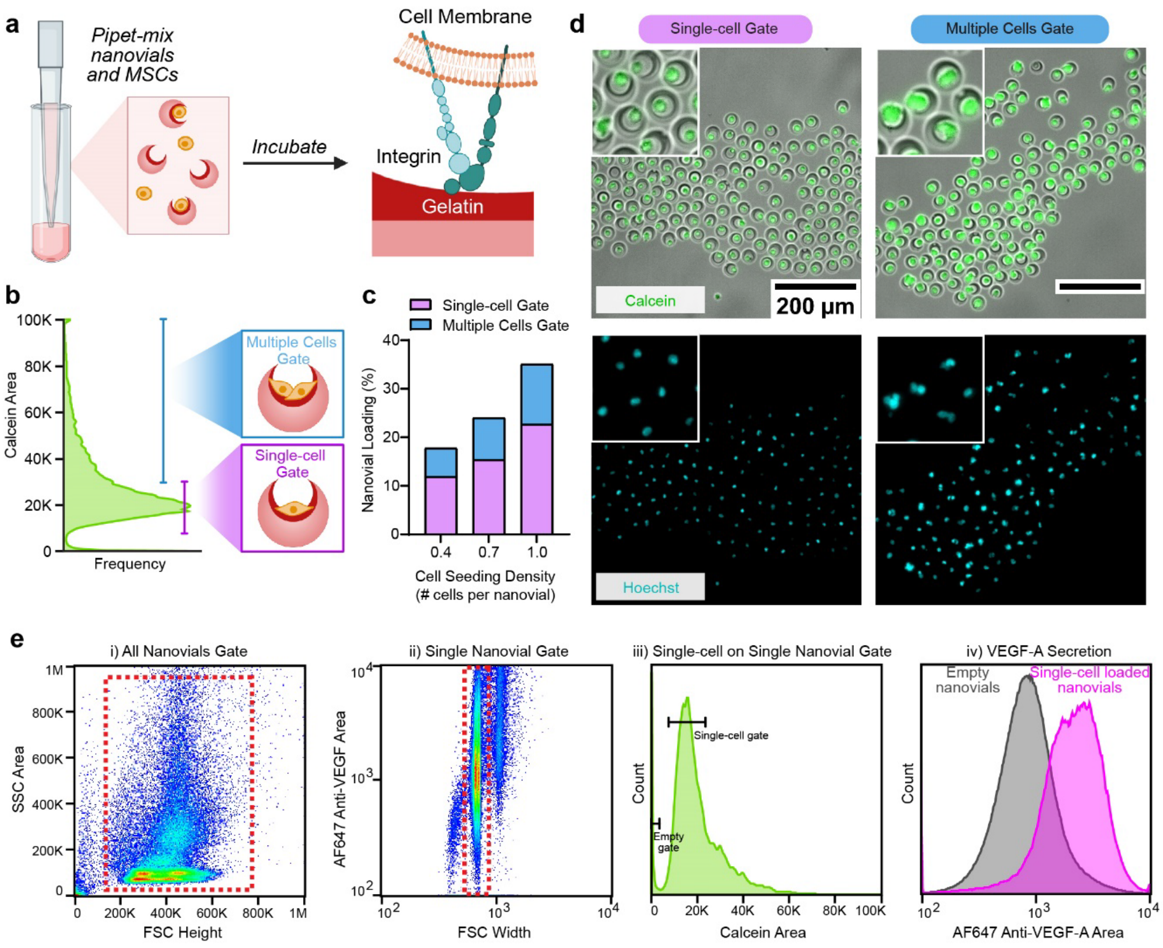


**Figure S1: Cell loading into nanovials, enrichment of single cell-loaded nanovials by FACS, and VEGF-A nanovial secretion assay validation. a,** Schematic showing anchorage of single cells on nanovials coated with gelatin via integrin binding. Cell loading into nanovials is achieved by simple mixing. **b,** Flow cytometry histogram of cell-loaded nanovials stained with calcein AM viability stain. Cells are sorted via FACS (SONY SH800S) based on calcein signal into ‘Multiple Cells’ and ‘Single-cell’ gates. The distribution of the calcein signal was found to have a peak with a tail at higher intensities, representing nanovials containing more than one cell. **c,** We tested three cell loading concentrations (0.4, 0.7, and 1 cell per nanovial) and analyzed the fraction of nanovials carrying single cells using the gates described in (b). Graph quantifying cell loading into nanovials, depicting the proportion of nanovials with one cell or multiple cells. When loading cells at the 1:1 cell-to-nanovial ratio, we achieved ~23% single-cell loaded nanovials which could be separated by sorting for downstream approaches and analyses. **d,** Fluorescence microscopy images of nanovials sorted for the indicated gates as described in (b). By sorting with nanovials in the ‘Single-cell’ gate, we enriched for nanovials carrying single cells as confirmed by Hoechst nuclei staining, whereas nanovials in the tail (‘Multiple Cells Gate’) represented mostly two or more loaded cells. Following sorting, we estimated that 95% of the “Single-cell” gate sorted nanovials contained one cell by image analysis. **e,** Gating strategy used for cell-loaded nanovial samples. i) All nanovials were gated in FSC/SSC, followed by ii) a single nanovial gate based on FSC-Width, and iii) the ‘Single-cell’ gate based on calcein signal intensity was used to isolate single cells on single nanovials. iv) Flow cytometry fluorescence histogram of single cell-loaded nanovials and empty nanovials after 12 hours of secretion incubation. Single cell-loaded nanovials have higher AF647 anti-VEGF-A signal than empty nanovials isolated from the same sample, showing low crosstalk.

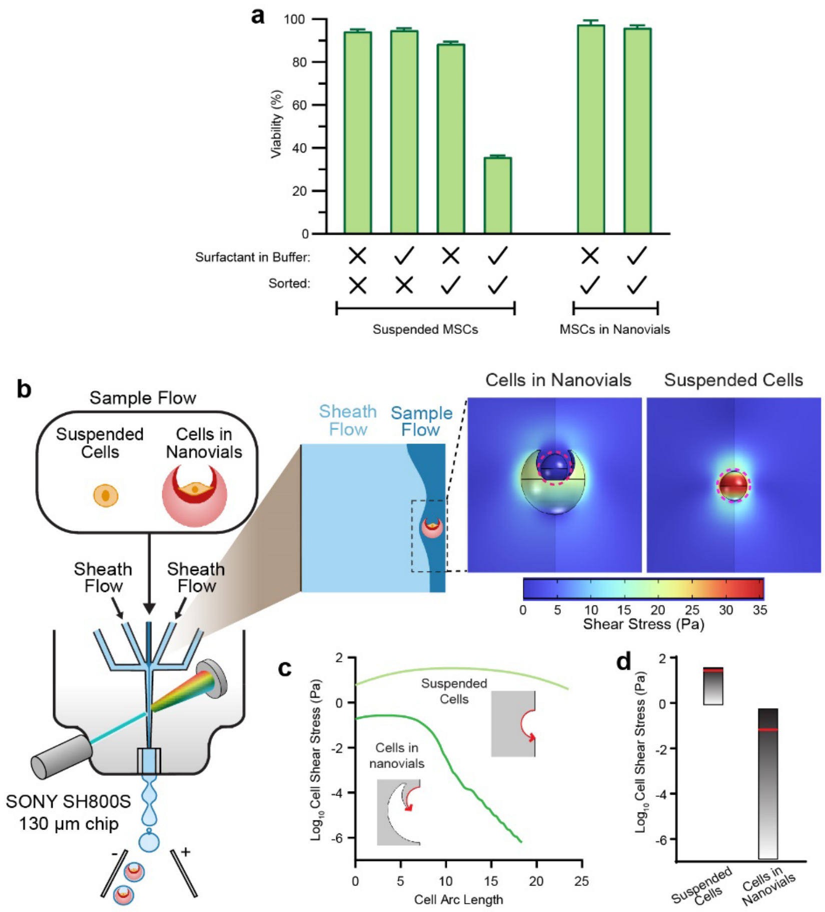


**Figure S2: Nanovials protect viability of MSCs during flow sorting. a,** The effect of surfactant and sorting on viability of cells on and off nanovials, as measured by live/dead stain imaging. Typically, nanovial samples are kept in buffer with a surfactant (Pluronic) at low concentration for handling and sorting steps, as it prevents nanovials from aggregating. Here, we exploit the surfactant as a stressor to test the effect of sorting in and out of nanovials on MSC viability. For suspended MSC samples, MSCs were dissociated from flasks, resuspended in FACS buffer with and without Pluronic surfactant, and viability was measured for MSCs with and without sorting. For MSC-loaded nanovial samples, MSCs were loaded on nanovials, resuspended in wash buffer with and without Pluronic surfactant, and viability was measured for MSCs after sorting. We found that viability decreased significantly when MSCs suspended in FACS buffer with Pluronic are sorted, but all other conditions maintained high viability. Surfactant, even at low concentrations, can likely induce some membrane damage, which is further damaged during sorting; however, nanovials seem to protect cells from this damage. **b,** Finite element modeling using COMSOL results show that cells in nanovials are exposed to reduced levels of shear stress compared to cells in suspension when flowing through the nozzle of a flow sorter (see methods). Here, the shear stress is plotted on the cell and nozzle geometry, and shows how the suspended cell (right) experiences greater shear stress than the cell inside nanovial (left). **c,** Shear stress from the COMSOL model is plotted against position along the cell perimeter for suspended cells and cells adhered within a nanovial. The red arrow in each schematic (based on the model geometry) indicates the direction of arc length and shear stress measurement. **d,** Range of shear stress for suspended cells and cells adhered within a nanovial based on (c), with average shear stress plotted (red line). The average shear stress is 400-fold higher for suspended cells than cells in nanovials.

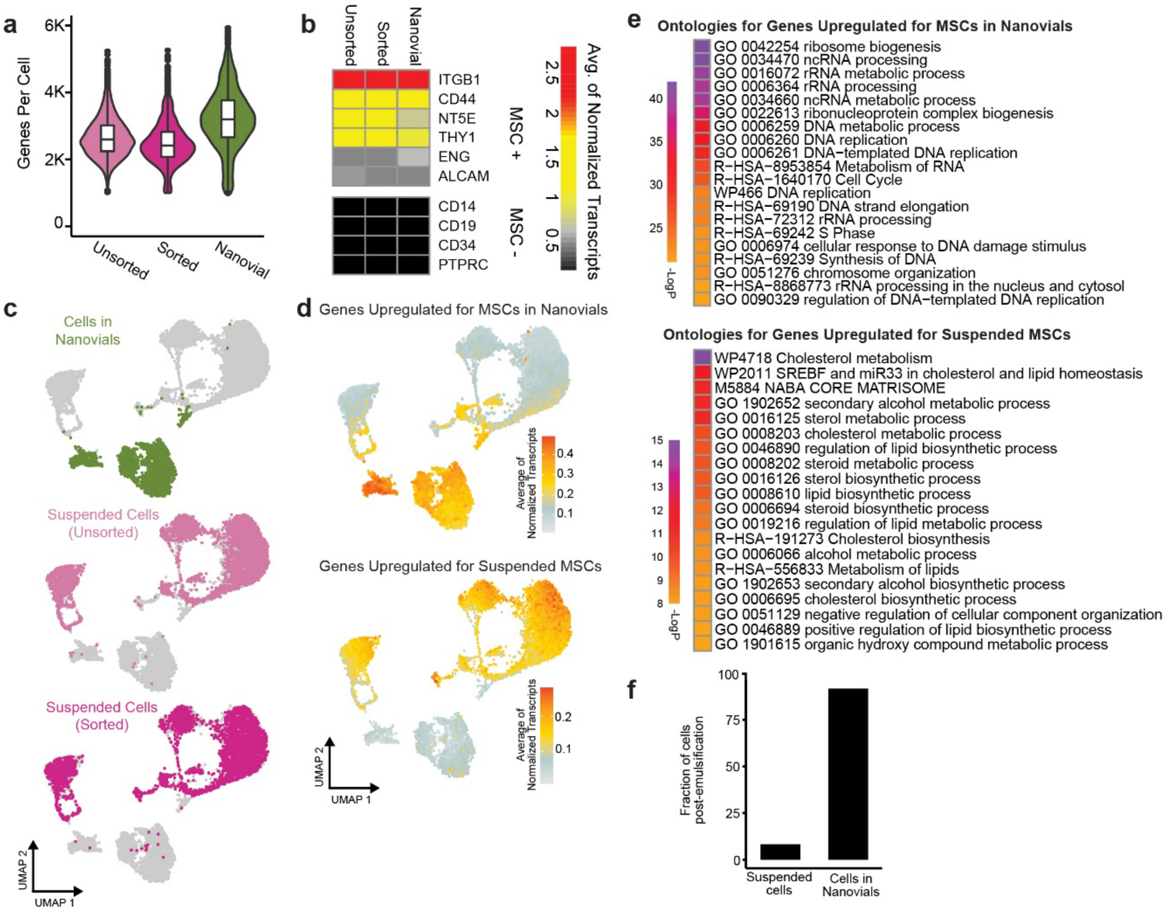


**Figure S3: Transcript changes related to cell loading and adhesion in nanovials. a,** For the scRNA-seq experiment with MSCs loaded into nanovials or freely suspended shown in Figure 2i, the graph shows the genes per cell for suspended and unsorted MSCs, suspended FACS-sorted MSCs, and MSCs loaded on nanovials and sorted. **b**, Heatmap showing the average normalized transcript levels of known MSC markers (top) and markers from other cell types (bottom) in each condition from the experiment in (a) (suspended, unsorted; suspended, sorted; on nanovials and sorted). **c**, UMAPs of the combined transcriptome data from the scRNA-seq experiment described in (a). The cells from each condition are separately displayed and colored. **d**, As in (c), showing the mean transcript level of genes significantly upregulated in cells adhered to nanovials relative to suspended MSCs (top) or upregulated in suspended MSCs (bottom). **e**, Gene ontology for the two gene sets from (d). **f**, Proportion of cells in nanovials that could not be associated with a nanovial-tagged oligo-barcode and are therefore presumably detached or can be associated with an oligo-barcode and therefore presumably are still on the nanovial after the emulsification step.

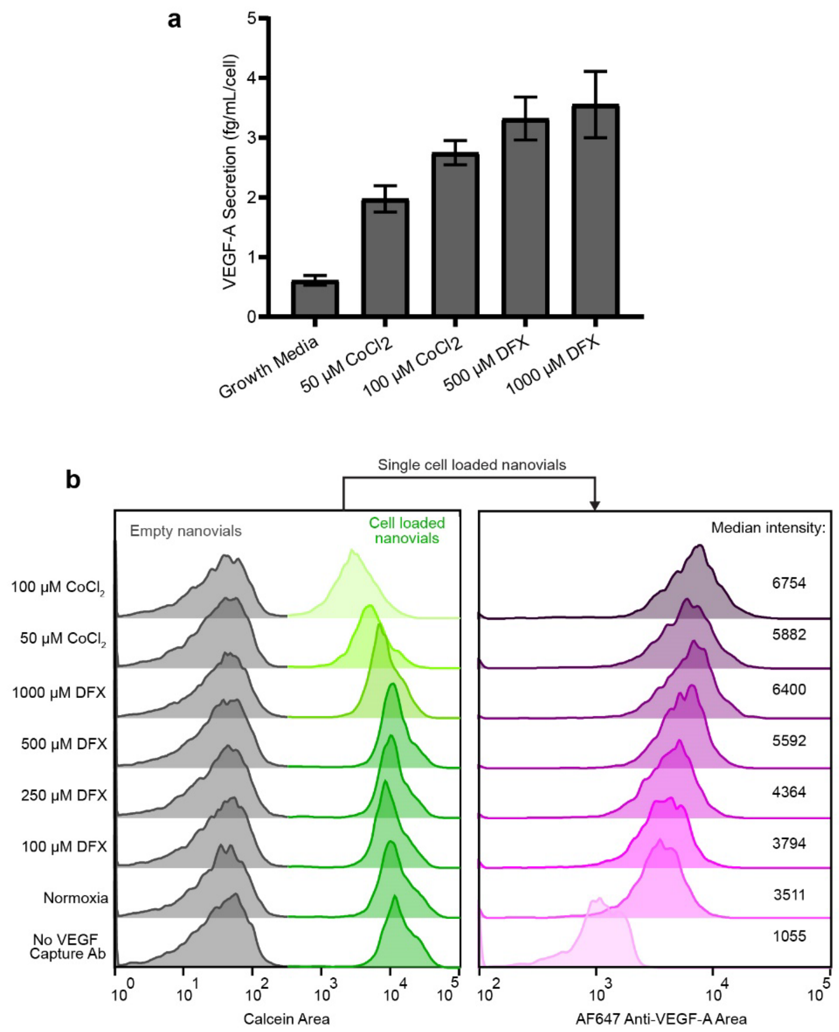


**Figure S4: Effect of hypoxia inducers on VEGF-A secretion by MSCs. a,** ELISA for VEGF-A secretion from conditioned media from MSCs grown on tissue culture plates under normoxic condition (normal growth media) and treated with indicated concentrations of cobalt chloride ($CoCl_2$) and deferoxamine (DFX) hypoxia mimicking agents for 24 hours to induce hypoxic conditions. **b,** Flow cytometry histograms are shown for two fluorescence channels indicating calcein positive MSC-loaded nanovials with anti-VEGF-A labeling on nanovials for MSCs treated with indicated concentrations of cobalt chloride ($CoCl_2$) and deferoxamine (DFX) hypoxia mimicking agents for 12 hours. Normoxia and no anti-VEGF-A capture antibody controls are also shown. 500 µM DFX yielded the largest increase in VEGF-A secretion (AF647 anti-VEGF signal) without compromising cell metabolic activity/viability (calcein).

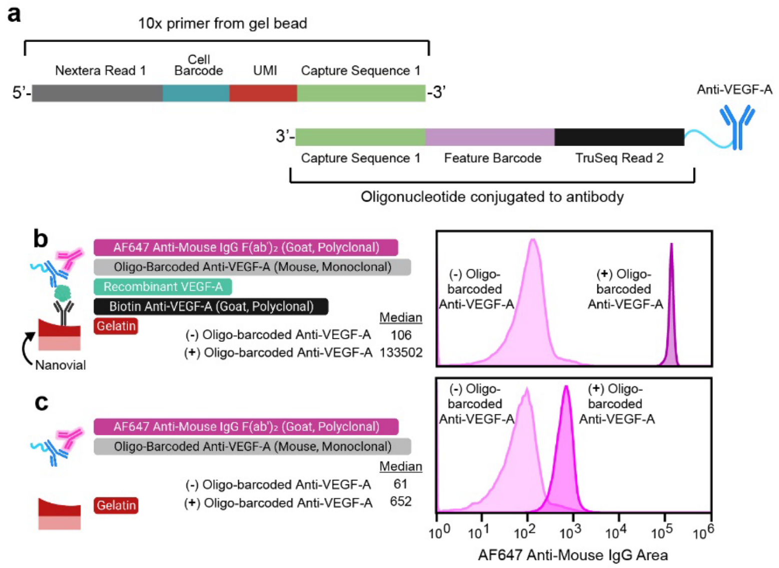


**Figure S5: Oligo-barcoded Anti-VEGF-A binding specificity on nanovials. a.** An anti-VEGF-A antibody (to be used as detection antibody for VEGF-A secretion in our SEC-seq approach) was conjugated with a 10X compatible oligo-barcode along with necessary sequences for 10x library preparation. The schematic shows the sequence composition of the oligo attached to the VEGF-A detection antibody, along with the 10x primer which hybridizes to the antibody-derived oligo and adds the unique molecular identifier (UMI) upon reverse transcription. **b,** (Left) Schematic showing the attachment of and the detection immunoassay for recombinant VEGF-A via the oligo-barcoded antibody described in (a) which was quantified with a fluorescently-labeled secondary antibody in nanovials by flow cytometry. (Right) Flow cytometry histograms showing outcome of the experiment on the left with recombinant VEGF-A with and without the oligo-barcoded VEGF-A detection antibody, demonstrating antibody specificity of the detection assay and the validity of the oligo-barcoded VEGF-A detection antibody to the presence of VEGF-A protein. The numbers on the left indicate the median of both histograms. **c**, As in (b), except that no biotinylated anti-VEGF-A capture antibody and recombinant VEGF-A was used in the assay.

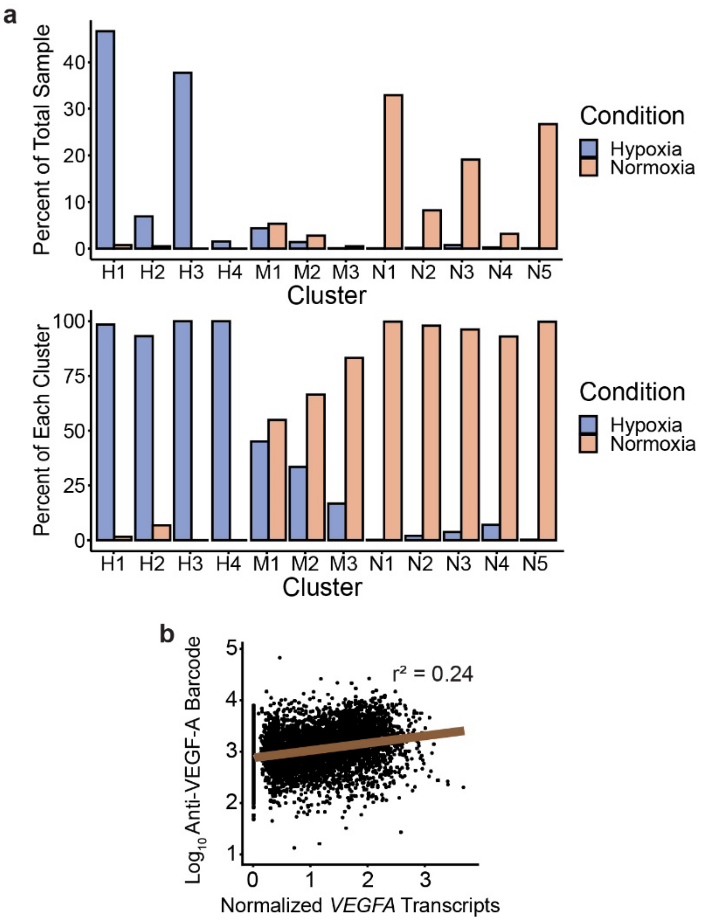


**Figure S6: Analysis of the SEC-seq experiments for normoxic and hypoxic MSCs. a,** Distribution of hypoxic and normoxic MSCs from the SEC-seq experiments described in Figure 3, per cluster depicted in Figure 3f as a percent of each sample or as a percent of each cluster. **b**, Scatter plot shows the correlation between *VEGFA* transcript and VEGF-A secretion for individual cells from both hypoxic and normoxic conditions.

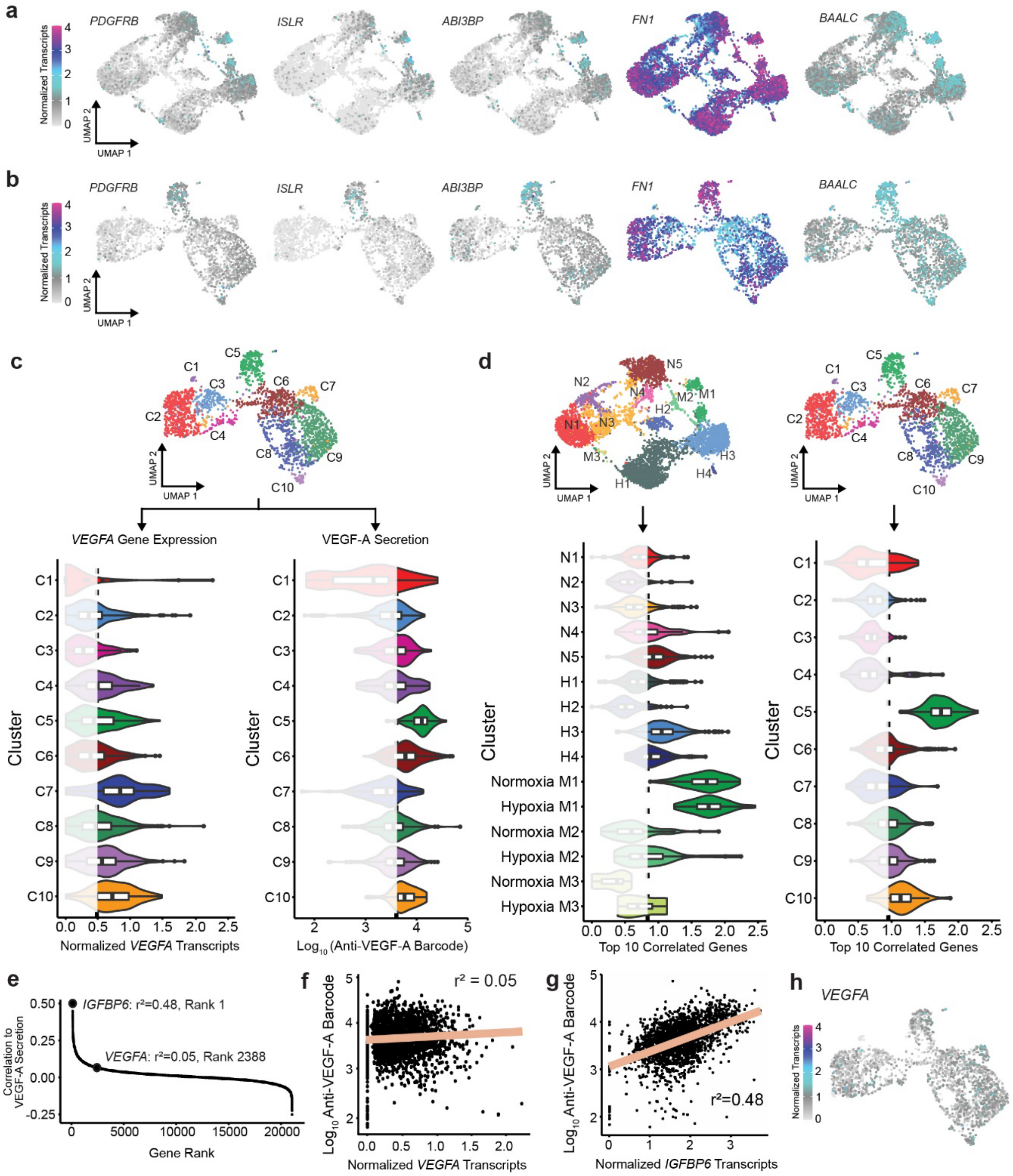


**Figure S7: Identification of a high-VEGF-A secreting MSC subpopulation in a replicate experiment. a.** UMAPs showing the normalized transcript level of the indicated genes for the normoxic/hypoxic SEC-seq experiments from Figure 3. All five genes belong to the top 10 correlates for VEGF-A secretion. **b.** As in (a), for the replicate normoxic SEC-seq experiment from Figures 4d/e **c,** Violin plots by cluster showing VEGF-A secretion and *VEGFA* transcript levels for all cells in the SEC-seq experiment for normoxic MSCs from (b), for the clusters shown in Figure 4e. **d,** Violin plots showing the average normalized transcript level of the 10 best correlating genes with VEGF-A secretion from Fig. 4b for the SEC-seq experiments with normoxic and hypoxic MSCs from Figure 3 and the normoxic replicate. **e,** For the SEC-seq experiment with normoxic MSCs in (b), all detected genes were ranked by the correlation of their transcript levels to the VEGF-A secretion level. Each gene is plotted by its rank and correlation. The ranks of the *VEGFA* and *IGFBP6* genes are highlighted and the correlation value is given. **f-g,** Scatter plots showing **f,** the correlation between *VEGFA* transcript and VEGF-A secretion for individual cells and **g,** the expression of IGFBP6 normalized transcripts vs the log transformed VEGF-A secretion reads in the replicate normoxic MSC experiment. Correlation value shown on plot and displayed via the linear regression line. **h,** UMAP showing *VEGFA* transcript levels per cell for the replicate normoxic SEC-seq experiment

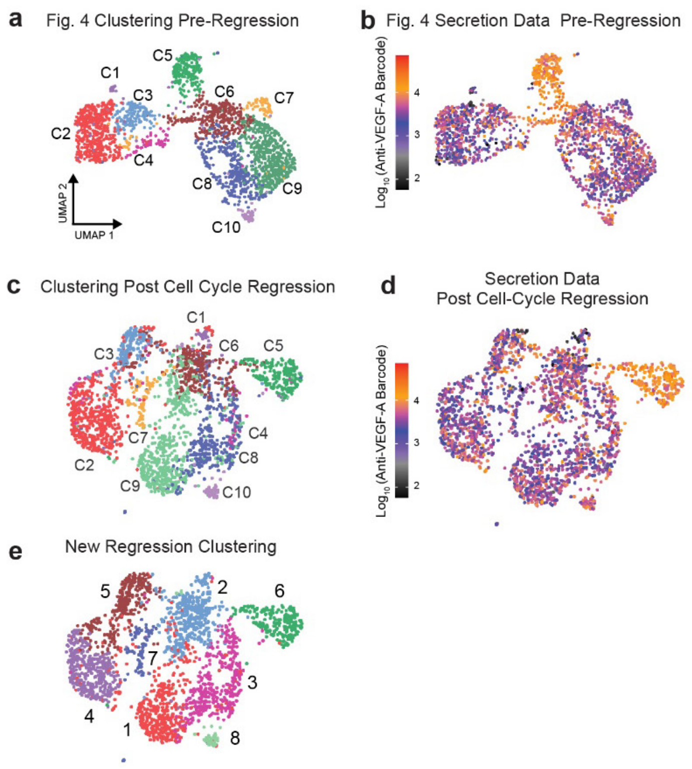


**Figure S8: The high VEGF-A secretion cluster is not affected by cell cycle regression. a**, UMAP of the cell clustering in the normoxic SEC-seq replicate from Figure 4 is shown again for comparison with the cell cycle-regressed data. **b**, VEGF-A Secretion per cell for the normoxic SEC-seq replicate from Figure 4 is also shown again for comparison with the cell cycle-regressed data. **c,** New UMAP coordinates and clustering of normoxic cells in (a) post cell cycle regression. Note that Cluster C5's spatial separation from other clusters is preserved with low mixing. **d,** VEGF-A secretion shown on the new UMAP coordinates post cell cycle regression. The cells in the newly arranged cluster C5 remain highly enriched for high VEGF-A secretion. **e,** New clustering of cells based on the cell cycle regression modified data is displayed on the UMAP. While the borders between other clusters has shifted, the majority of cells that made up cluster C5 still distinctly form their own cluster, #6, demonstrating that the highly secretion cluster's special transcriptional profile is unaffected by cell cycle regression.
